# Stochasticity in Metabolism: Oxidative Stress Response Phenotypes As Emergent Consequences of Network Topology

**DOI:** 10.64898/2026.10.23.761362

**Authors:** Alex Mihnev, Dana Luong, Jonah Porat, Roger Chang

## Abstract

Reactive oxygen species (ROS) are ubiquitous yet elusive due to their reactivity and short half-lives, making simulation essential for mechanistic insight. Traditional ROS modeling has been confined to small ODE frameworks limited by kinetic-parameter scarcity. Here, we recast ROS dynamics as a flux-balance analysis (FBA) optimization problem using COBRA and CPLEX, augmenting the *Escherichia coli* iJO1366 genome-scale model with 84 spontaneous oxidation, 14 redox, and 9 detoxification reactions, targeting three ROS species: hydroxyl radicals (•OH), superoxide (•O_2_-), and hydrogen peroxide (H_2_O_2_). We also formalize five fundamental rules of oxidation biology in the form of mixed-integer linear constraints, allowing us to dynamically incorporate reaction rate constants and let complex behavior emerge from first principles. Among these phenomena is the *bimodal killing pattern* under H_2_O_2_ stress, first announced over 40 years ago and never computationally recapitulated until now. We observe it by stochastically sampling our models with GapSplit, simulating bet-hedging and partitioning solutions into growers, persisters, and non-survivors. In our simulations, this killing effect arises solely from the metabolic network topology during random exploration of alternative metabolic configurations. The conservation of steady state creates a stochastic bias for detoxification when central carbon metabolism reaches its metabolic capacity to deal with ROS, causing the famous survival recovery phase. The recovery phase is thus part of Mode 1 metabolism, characterized by spare detoxification capacity and fulfilled by the Ahp reaction in *E. coli*. Mode 2 has no spare detoxification. We anticipate this to be an overlooked universal principle.

## Introduction

Reactive Oxygen Species (ROS) are a ubiquitous biological stress. The fundamentality of ROS as a stress is highlighted by anaerobic organisms also experiencing and having to defend against it (Bao et al., 2017; Imlay, 2019; Liu et al., 2011; Lu & Imlay, 2021; Tong et al., 2007; Wang et al., 2019). Although nominal ROS production is a part of normal function of many cellular systems (Milkovic et al., 2019), high ROS is commonly associated with damage to macromolecules (Davalli et al., 2018; Juan et al., 2021; Sies et al., 2024) such as proteins (Amici et al., 1989; Bollineni et al., 2011; Cheignon et al., 2018; Garrison, 1987; Hawkins & Davies, 2019; Maisonneuve et al., 2009; Matallana-Surget et al., 2013; Mirzaei & Regnier, 2006; Stadtman & Levine, 2003; Taborsky & McCollum, 1973; Temple et al., 2006), membranes and lipids (Ito et al., 2019; Kao et al., 2020; Yadav et al., 2019), DNA (Lawless et al., 2020; Poetsch, 2020; Salehi et al., 2018; Srinivas et al., 2019; Yan & Zaher, 2019; Zheng et al., 2001), and iron-sulfur clusters (Imlay, 2006; Jastroch et al., 2010; Read et al., 2021; Santos-Sánchez et al., 2019; L. Yang et al., 2019). Spontaneous oxidation reactions (SORs) damage these biomolecules and cause a cascading cellular system failure, which can contribute to disease development such as cancer (Baffy, 2010; Green et al., 2024; Milkovic et al., 2019; Srinivas et al., 2019; Zhang et al., 2006), cardiovascular disease (Kao et al., 2020; Taverne et al., 2013), neurodegenerative conditions (Cheignon et al., 2018; Cioffi et al., 2019; Höhn et al., 2020; Kao et al., 2020), and general effects of aging (Franceschi et al., 2018). A unifying feature of these pathologies in eukaryotes is mitochondrial dysfunction associated with elevated ROS (Diaz-Vegas et al., 2020; Franceschi et al., 2018; Lawless et al., 2020; Yuan et al., 2020) while co-occurring with other oxidation markers like protein aggregation and lipid peroxidation (Cheignon et al., 2018; Gianazza et al., 2021; Kao et al., 2020; B. Yang et al., 2019) dityrosine crosslinking, methionine oxidation, 4-hydroxinonenal accumulation, heme depletion, tau enriched neurofibrillary tangles (Cioffi et al., 2019) 8-oxo-7,8-dihydroguanosine (Fasnacht & Polacek, 2021), and others. Studying ROS metabolism in tractable model systems like *Escherichia coli* could unlock actionable insight for pathogens, diseases, and treatments. The flow of ROS through the metabolome is difficult to measure directly (Takhaveev & Heinemann, 2018), instead relying on persistent secondary effects of polymer modifications. Despite accounting for only 2% of the cell’s dry weight (Neidhardt et al., 1990), the metabolome provides the energy and matter required to support every process in the cell. Therefore, network-wide simulation of ROS metabolism potentiates broad fundamental insight about mechanisms of oxidative stress, including events which would otherwise be impossible to detect. Accounting for the direct effects of ROS on small molecule metabolites is enabled by broad knowledge of the chemistry of SORs (Bielski et al., 1985; Buxton et al., 1988) incorporation of this knowledge into a genome-scale metabolic model (GEM) permits simulation of oxidative stress to study how it effects the distribution of metabolic flux and how this may affect cellular phenotypes.

### Sources of ROS

Three primary ROS are most important in the context of cellular oxidative stress: hydroxyl radical (•OH), superoxide (•O_2_-), and hydrogen peroxide (H_2_O_2_) (Krumova & Cosa, 2016). Normally generated through the incomplete reduction of molecular oxygen (Imlay, 2013), the major cellular sources of ROS are as side products of the electron transport chain (ETC) (Collin, 2019; Mailloux, 2020; Nissanka & Moraes, 2018) and certain redox enzymes (Addorisio et al., 2022; Ambrus et al., 2011; Geary & Meister, 1977; Imlay, 1995; Korshunov & Imlay, 2010; Kumar & Imlay, 2013; Kussmaul & Hirst, 2006; Massey et al., 1969; Messner & Imlay, 1999; 2002; Snezhkina et al., 2019; Szabó & Ambrus, 2023) – mostly in the form of H_2_O_2_ and •O_2_-. Light and other forms of radiation also produce additional ROS types (Asada, 1999; Forenzo & Larsen, 2024; Garrison, 1987; Guo et al., 2024; Wang et al., 2019) but these are not in the scope of this work. Oxidative stress in organisms like *E. coli* may also be naturally produced in the growth medium (Li & Imlay, 2018), by host defenses (Collin, 2019; Dahlgren et al., 2019; Lam et al., 2020; Magnani & Mattevi, 2019; Matilla & Krell, 2018; Nguyen et al., 2017; Segal, 2008; Turrens et al., 1985; Winterbourn, 2013), pharmaceuticals (Turrens, 2003), or invoked through exogenously provided H_2_O_2_ which, unlike superoxide, can cross the cell membrane (Fasnacht & Polacek, 2021; Seaver & Imlay, 2001a). H_2_O_2_ does not generally cause direct molecular damage, and superoxide does not damage DNA (Winterbourn, 2013). Instead, both participate in the metal-catalyzed Fenton reaction and Haber-Weiss cycle (Chen, 2019; Fenton, 1894; Imlay, 2006; Kehrer, 2000; Koppenol, 2001; Liochev & Fridovich, 2002), generating •OH, which is the primary biomolecule-damaging ROS (Winterbourn, 2013). H_2_O_2_ is a key for SOR activity because it can cross biological membranes and be directly converted to •OH. This can explain why this peroxide, also generated through normal metabolic activity, is carefully handled by the cell. Metabolism maintains the steady state concentration of H_2_O_2_ to around 50 nM by detoxification systems (Imlay, 2013) while transcriptional responses, such as those initiated by the OxyR regulon, react to as little as 0.1 µM extracellular H_2_O_2_ (Seaver & Imlay, 2001a). Most of these detoxification systems have evolved to clear H_2_O_2_ directly, thus preventing damage.

### ROS detoxification systems

Alkyl hydroperoxide reductase (Ahp), catalyzing the NADHPO reaction, is the principal detox system indispensable for fitness under continued oxidative stress (Seaver & Imlay, 2001b). Unlike most other detox reactions that use NADPH for reducing potential, Ahp uses NADH, which could explain the conditional synergy with the two catalases which use neither (Seaver & Imlay, 2001b). The two catalases, HPI and HPII, have a high capacity for H_2_O_2_ but lower affinity than Ahp and are only expressed in response to exposure to H_2_O_2_ or in stationary phase (Loewen et al., 1985). Catalases are responsible for most detox at higher concentrations of H_2_O_2_, whereas Ahp dominates at lower (Uhl & Dukan, 2016). The MsrA and MsrB detox systems (Moskovitz, 2005; Tarrago et al., 2012) use NADPH, targeting methionine-S- and -R-sulfoxides, respectively. Msr detoxification works downstream of molecular damage rather than targeting H_2_O_2_ directly. Additional ROS detox systems include peroxiredoxins (Imlay, 2008), superoxide dismutases (SOD) and spontaneous dismutation (SD) of superoxide (Palma et al., 2020; Wen & Garg, 2018), thioredoxins (Lin, 1999) like the thioredoxin-dependent H_2_O_2_ reductase (Trx-HPR), and the glutathione peroxidase (GPx) reaction (Mishra & Imlay, 2012). However, enzymatic detoxification is energetically expensive due to consuming reducing equivalents, meaning that the most efficient defenses could be mechanisms to avoid excessive ROS exposure, also known as *persistence*.

### Stochastic survival strategies: growth and persistence

Persistence is a stochastic effect in which subpopulations of cells spontaneously change in response strategies under stress, where some grow actively (growers) while others (persisters) go dormant to reduce susceptibility to ROS or other stresses and increase chronological lifespan which increases the odds of outliving the stress. The stochasticity of survival strategies is typically attributed to bacterial monocultures (Raser & O’Shea, 2005; Tonn et al., 2019; Wehrens et al., 2018) but is not exclusive to bacteria and has also been observed in yeast (Shabestary et al., 2024) and diatoms (Mizrachi et al., 2019). A relevant example of persistence is H_2_O_2_ exposure causing a prolonged lag phase or dormancy in *E. coli*, followed by otherwise normal exponential growth (Haque et al., 2025; Jozefczuk et al., 2010; Roth et al., 2022; Sahu & Behuria, 2018) which may be caused by translational inhibition due to oxidative stress (Zhong et al., 2015; Zhu & Dai, 2019). The duration of dormancy is generally determined by the amount that a toxin such as H_2_O_2_ exceeds a cellular threshold (Rotem et al., 2010). This suggests that most *E. coli* cells enter dormancy as persisters and only resume growth once ROS has been sufficiently reduced either spontaneously or though persistent detox systems.

### Bethedging and metabolic hysteresis

Some individuals will naturally die when exposed to a stress, but others will survive. Those that survive, be it growers or persisters, are pre-determined by the pre-exposure phenotypic heterogeneity of the population which, when the exposure is short enough, carries over to the stressed condition through a form of metabolic inertia, called *hysteresis* (Oyarzún & Chaves, 2015). This strategy involving natural subpopulation heterogeneity is known as *bet-hedging* (Bertrand, 2019; Veening et al., 2008) and enables species survival of sudden-onset stresses. The effect is stochastically influenced so that a proportion of cellular metabolic phenotypes predisposed towards survival of a specific stress may have evolved in response to the probability of experiencing that stress (Kussell & Leibler, 2005).

The dormancy of *E. coli* under H_2_O_2_ stress is well-explained by this theory, given the selective pressure posed by exposure of the bacterium to host immune systems. Having a translational apparatus sensitive to ROS (Zhong et al., 2015; Zhu & Dai, 2019), ensuring an overwhelming majority of persisters, would thus be an asset and not a liability.

### Bimodal fitness patterns

A relevant effect that simulations may be able to explain is the fraction of survivors as a function of time or stress – both of which are non-monotonic functions. Bet-hedging explains what individuals survive but the number of survivors seems to change non-linearly with the passage of time or extent of stress. This implies the onset of metabolic context shifts which affect what sub-groups of individuals, be they growers or persisters, are more likely to survive. Most examples in the literature consist of a steep linear rate of death, as a function of time after stress, followed by shifting to a more shallow rate past a certain point – known as the *bimodal survival pattern* (Balaban et al., 2004; Boldrin et al., 2020; Brauner et al., 2016; Fanous et al., 2025; Gefen & Balaban, 2009; Quigley & Lewis, 2022; Rotem et al., 2010; Shultis et al., 2022). Another way to analyze survival fractions is as a *bimodal killing pattern*, not to be confused with the more mentioned bimodal survival, where treatments are dispensed for the same length of time, usually on the order of minutes, and survivors are recovered and counted on solid rich media. Killing is then a function of treatment intensity.

The bimodal killing pattern of *E. coli* in response to H_2_O_2_ was described in the 1980s (Brandi et al., 1987; Imlay & Linn, 1986). It consists of an initial steep decline in survival (Mode 1 killing), followed by a gradual recovery, and terminating in a long and final dropoff in survival (Mode 2 killing). All of this is a function of increasing H_2_O_2_ concentration. This killing pattern has been observed between 0 and 20 mM H_2_O_2_, where Mode 2 killing begins around 10 mM H_2_O_2_. The absolute survival limit of *E. coli* is around 50 mM H_2_O_2_ (Zhu & Dai, 2019) and is achieved by pre-exposure to lower concentrations of H_2_O_2_ inducing catalase expression (Rodríguez-Rojas et al., 2020). The mechanism behind Mode 1 killing has been debated but is generally considered to be linked to DNA damage, whereas Mode 2 killing has been associated with limits to polymer stability, notably cell membranes, upon oxidation (Asad et al., 2004; Brandi et al., 1987; Linley et al., 2012). Despite these explanations, the bimodal killing pattern has been notoriously difficult to model mathematically, implying that there may be important overlooked contributions from metabolism. A prior study (Uhl et al., 2015) presented an Ordinary Differential Equations (ODEs) model predicting a decrease in iron availability associated with Mode 1 killing, with a clear connection to cell size. This implicated an important metabolic component involved in ROS production and the ability to dilute damage by increasing cell volume.

Bimodal distributions are known to be formed stochastically (To & Maheshri, 2010) and can be stabilized by positive feedback loops (Ge et al., 2018). Understanding the mechanism behind the bimodal killing pattern could be critical in medicine as well as systems biology as this effect may be more ubiquitous than previously thought and potentially impact cells and mitochondria in disease. For example, this effect has also been observed in Chinese hamster ovary cells exposed to H_2_O_2_ (Kaneko et al., 1994).

### Previous strategies for simulating ROS metabolism

Considering everything we know about ROS detoxification, heterogeneity of cellular states, persistence, hysteresis, and bimodal distributions, we propose that a stochastic model of metabolism could explain the bimodal killing pattern and provide fundamental biological insight into oxidative stress phenotypes. Most previous attempts to simulate systemic effects of ROS on the metabolism of *E. coli* have relied on ordinary differential equations (ODEs) (Schleicher, 2021). Importantly, ODEs are usually deterministic and always kinetic, capturing the temporal dimension but requiring rate parameters for each reaction to achieve biological realism. Experimentally determined kinetic parameters are not available for most *E. coli* metabolic enzymes, resulting in smaller, less comprehensive theoretical models for ROS simulations (Uhl et al., 2015). Such an approach does not suffice to investigate the stochastic nature of ROS metabolism under stress, nor does it lend itself to more comprehensive models.

Constraint-based modeling using flux balance analysis (FBA) (Orth et al., 2010), commonly associated with the COBRA toolbox (Heirendt et al., 2019), is a linear programming (LP) graph optimization approach for metabolic network simulations. This technique scales exceptionally well due to abstracting time and rate parameters with a biomass objective function at steady state and has even been applied to simulate the systemic effects of ROS (Bhalla et al., 2022; Monk et al., 2017; Sridhar et al., 2023; L. Yang et al., 2019) however, such studies have been limited in scope and particularly have not properly accounted for the realistic and dynamic distribution of spontaneous ROS reaction fluxes, which should not be expected to yield to maximizing an (growth) objective function. Nor have these studies accounted for the stochasticity of cellular metabolism.

### Stochastically sampled flux simulations

We present an alternative stochastic flux sampling approach which imitates bet-hedging in the context of ROS. This approach also formalizes five fundamental rules of ROS realism taking into consideration kinetic rate constants, enforced through dynamic constraints. We apply these rules, alongside expanded ROS topology, to the famous GEM of *E. coli* – iJO1366 (Orth et al., 2011). Our method is uniquely suited for studying short-timescale effects of oxidative stress, where observed effects arise naturally as a consequence of our rules and recapitulate several properties of *in vitro* cell cultures.

Our simulations effectively span the solution space for the model for each treatment while accounting for metabolic hysteresis from the pre-exposed state (Oyarzún & Chaves, 2015). Results from our simulated cell cultures exhibit stochastic properties in aggregate that offer new mechanistic insight to explain previously reported observations from *in vitro* studies, unifying much of the collective theory surrounding oxidative stress phenotypes in *E. coli* through a purely metabolic model of ROS. Thus, the realistic network-wide redistribution of fluxes upon oxidative stress may offer a simple explanation to some common effects of ROS, such as the bimodal killing pattern, which can translate to advancements in other areas of life science, in addition to providing fundamental insight.

### Modeling approach

A constraint-based modeling approach to study the relationship between ROS metabolism and cell survival in culture requires devising strategies to address several challenges, including: 1) broad representation of sources of ROS, spontaneous ROS chemistry, and ROS detoxification mechanisms; 2) enabling cells to adopt stochastic, heterogeneous metabolic states; 3) constraints to represent relative reaction rates of competing spontaneous oxidation reactions (SORs); and 4) evaluation of relative model accuracy to prioritize findings based on experimental metabolomics data. A major advantage of this approach is that the metabolic heterogeneity of individual cells can be represented by spanning the network-wide flux solution space through sampling. These sampled flux states can then be filtered based on specific behaviors, such as biomass flux to represent growth or distance to a reference state to represent hysteresis. We refer to our approach as an in silico cell culture, which could be performed using any available genome-scale metabolic network model (GEM). Below we describe how our modeling approach accommodates these intricacies of ROS metabolism through implementation of fundamental rules and demonstrate that this approach can recapitulate experimentally observed behaviors at the culture scale and provide detailed explanation of underlying metabolic mechanisms.

## Results

### Stochasticity, ROS detoxification, and steady state conservation drive bimodal killing

Stochasticity is a known cause of non-monotonic effects even in linear problems. We tested whether the bimodal killing pattern has a metabolic component by constructing a simplified topology of ROS metabolism operating through bet-hedging. The chosen topology of seven reactions represents central- and alternative carbon metabolism, spontaneous oxidation-, and ROS-detoxifying reactions (**Figure 1A**). The conservation of steady state is not only a constraint in all our simulations but also a thermodynamic property of real cells (Wilson & Matschinsky, 2021). We consequently hypothesized that metabolism explores alternative flux configurations to preserve steady state during increasing stress, manifesting in non-monotonic changes in fitness/ survival. Whenever these options are exhausted, we reasoned, a monotonic drop of fitness would represent the overwhelmed network. If such a network is unable to move ROS fast enough then it would no longer be a metabolic problem but a cascading system failure which is not within our scope.

**FIGURE 1.**
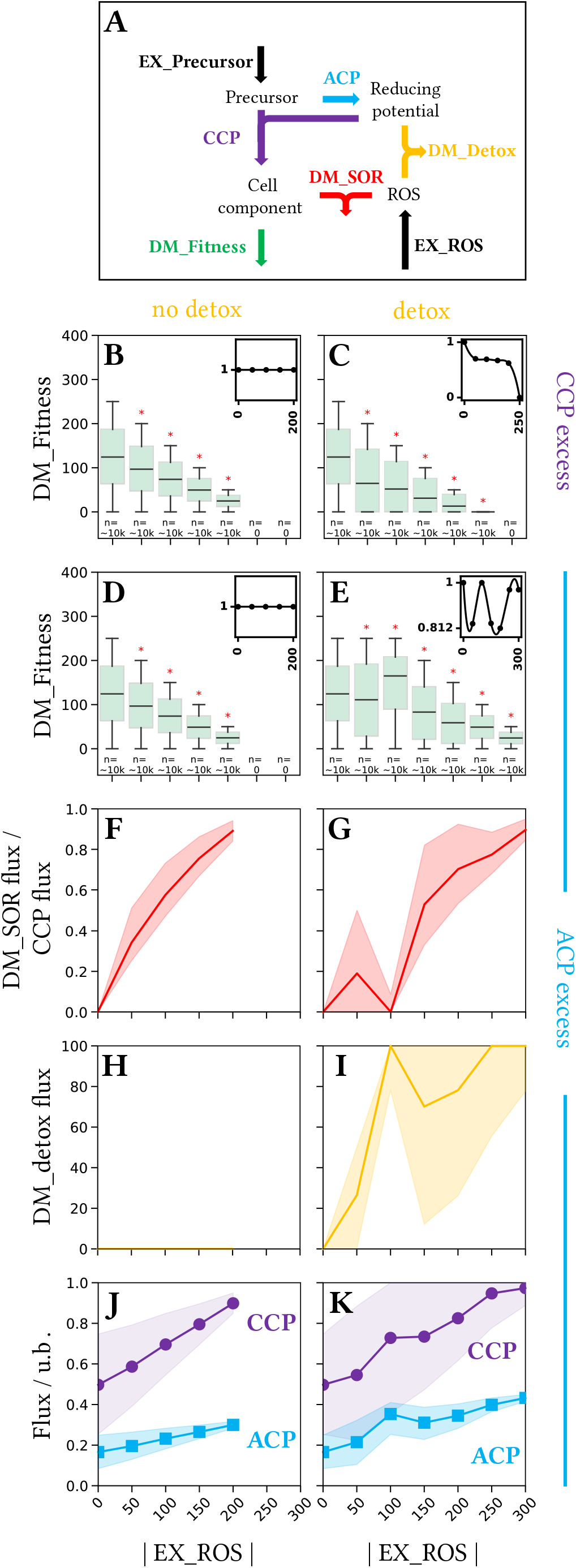
Bimodal killing can arise from a simple, ubiquitous metabolic topology and stochasticity alone. **(A)** Simple representation of steady state metabolism as a network. Four metabolites: Precursor, Reducing Potential, Cell Components, and ROS. Reactions: central carbon pathways (CCP, purple), alternative carbon pathways (ACP, blue), spontaneous oxidation reactions (SOR, red), and exchange reactions (EX_, black) for Precursor and ROS, DM_Fitness (green), DM_Detox (orange). **(B, C)** Fitness distributions without/with detox when CCP capacity exceeds ACP’s; insets show fraction of samples with net-positive fitness. Significance by t-test with Bonferroni correction: ns, p≥0.05; *, p<0.05. **(D, E)** Same, with ACP capacity exceeding CCP’s. Subsequent panels use ACP-excess condition only. (**F, G)** Median SOR flux as a fraction of CCP flux without/with detox; shading spans IQR. **(H, I)** Median DM_Detox flux with IQR. **(J, K)** Median CCP (purple) and ACP (blue), scaled to their upper bounds, with IQR.

Sampling of this model done stochastically with GapSplit (Keaty & Jensen, 2020) revealed no bimodal killing, regardless of detoxification, in conditions where central carbon metabolism has a higher capacity than alternative carbon pathways (**Figure 1B,C**) which generate reducing potential. Detoxification without an excess capacity for alternative reactions was only able to increase minimum fitness (**Figure 1C**). A bimodal killing pattern was instead seen when the alternative carbon pathways have a higher flux capacity and ability to detoxify (**Figure 1D-E**). The bimodal effect seemed to extend survival into higher concentrations of ROS via a recovery phase (**Figure 1E**). We defined Mode 1 as the metabolic configuration from increasing ROS manifesting in an initial drop in fitness (Mode 1 killing), leading up to and including the recovery phase. Mode 2 is associated with the following monotonic drop of fitness. Detox-incapable metabolism operates exclusively in Mode 2, as judged by SOR usage (**Figure 1F-G**). The detox reaction usage had a non-monotonic behavior, whenever present (**Figure 1H-I**). Under Mode 1 the network moves ROS indiscriminately through the SOR (**Figure 1G**) and/or detox (**Figure 1-I**) which was evident from the large variation of their usages. The blind increase of SOR usage thus explained the initial drop of fitness. Without detoxification, the activity of the central and alternative pathways increases linearly with ROS (**Figure 1J**). Activity through both reactions increases evenly with detoxification, followed by a more disproportionate utilization of central carbon metabolism’s full capacity (**Figure 1K**). This saturation of central carbon metabolism is due to its lower capacity combined with higher ROS. The only remaining way for steady state to move ROS is then through detoxification, creating a stochastic bias and a recovery phase. Subsequent exhaustion of this alternative carbon/detox path would mark the beginning of Mode 2. The prime requirement for recapitulating the bimodal killing effect is then:

- **Rule 1**: ROS-producing, consuming (SORs), and detoxifying reactions must be represented in the network topology

Given how ubiquitous this required topology is we anticipated that the bimodal killing pattern can be qualitatively validated in larger GEMs.

### ROS topology alone cannot reproduce bimodal killing in *E. coli*

For testing our findings in larger GEMs, we used a modified version of iJO1366 (Orth et al., 2011) as our Base model (see Methods). Among the changes to this model is the addition of a *death* constraint, preserving mathematical feasibility of solutions unable to handle oxidative stress, formalized as:

- **Rule 2**: the model can ignore external oxidative stress (H_2_O_2_) only if ATP maintenance flux is essentially zero

This constraint converts the models to a MILP, does not make dead solutions mechanistically interpretable, but – most importantly – allows direct counting of non-survivors. Furthermore, even well studied organisms like *E. coli*, where bimodal killing was observed, lack GEMs with properly represented spontaneous oxidation topology. This prompted us to systematically reconstruct 84 SORs, 9 detox, and 14 ROS-generating using only data with paired reaction rates (see Methods). The resulting *Base + Topology* (BT) model was subjected to an emulation of the conditions from Brandi et al. (1987) in growth-limited M9 medium (see Methods), paired with the Base model. Both models were thus appropriately dosed with H_2_O_2_, stochastically sampled with GapSplit, and corrected for hysteresis to simulate the short experiment timescale (see Methods). The results, shown in **Figure S1**, reveal no bimodal killing pattern of the surviving fraction.

Forcing the influx of H_2_O_2_ and losing metabolites to SORs must be a challenge for steady state. We anticipate that larger GEMs, like iJO1366, may use any means necessary to avoid ROS and many of which not possible in real life. This necessitates additional constraints.

### Dynamic rules of biochemically accurate simulation of ROS

Effects are most credible when they emerge from first principles rather than fitting data. The challenge was to then formalize a minimum set of first principles as non-spatiotemporal mathematical constraints (as fluxes are steady state rates of change, not dynamic time derivatives) to guide what means of handling ROS are permissible. What are the first principles of spontaneous oxidative biology then?

For starters, SORs are considered *instantaneous*, with higher *k* rate constants than enzymatic reactions. They should then take precedence and be obligatory as long as they are possible (i.e. both reactants are present):

- **Rule 3**: SORs must carry flux when all their reactants are present. Because fluxes are themselves rates, SOR (and detox) activity should be proportional to their *k* rates. This proportionality cannot be dynamically preserved by adjusting reaction bounds, at least not without severely debilitating the model:
- **Rule 4**: All SOR and detoxification fluxes must be a dynamic function of their respective reaction *k* constants multiplied with a scaling variable. It is tempting to want such scaling to be done with a single global variable. However, some SORs may be able to scale much higher than others who are limited by production of a reactant. Using a single variable would then undersample more extreme SOR differences. We thus need to use separate scaling variables for each reactant, expecting them to normally be coupled. Conditionally uncoupling these scaling variables must come with a penalty to prevent solver cheating:
- **Rule 5**: SORs/detox reactions sharing the same reactant may be scaled independently from all other SORs, provided that together they consume most of the affected reactant.

These first principles enforce and increase activity of rate-constrained reactions in lockstep. Additional oxidative stress would keep some of these reactions at their maximum while others may continue scaling up in lockstep, and so on until the death constrained has to be triggered. Most of these first principles are summarized in **Figure 2** while all are explained in detail in the Methods. It is important to note that detox reactions obey most of these first principles, but their usage is not compelled.

**FIGURE 2.**
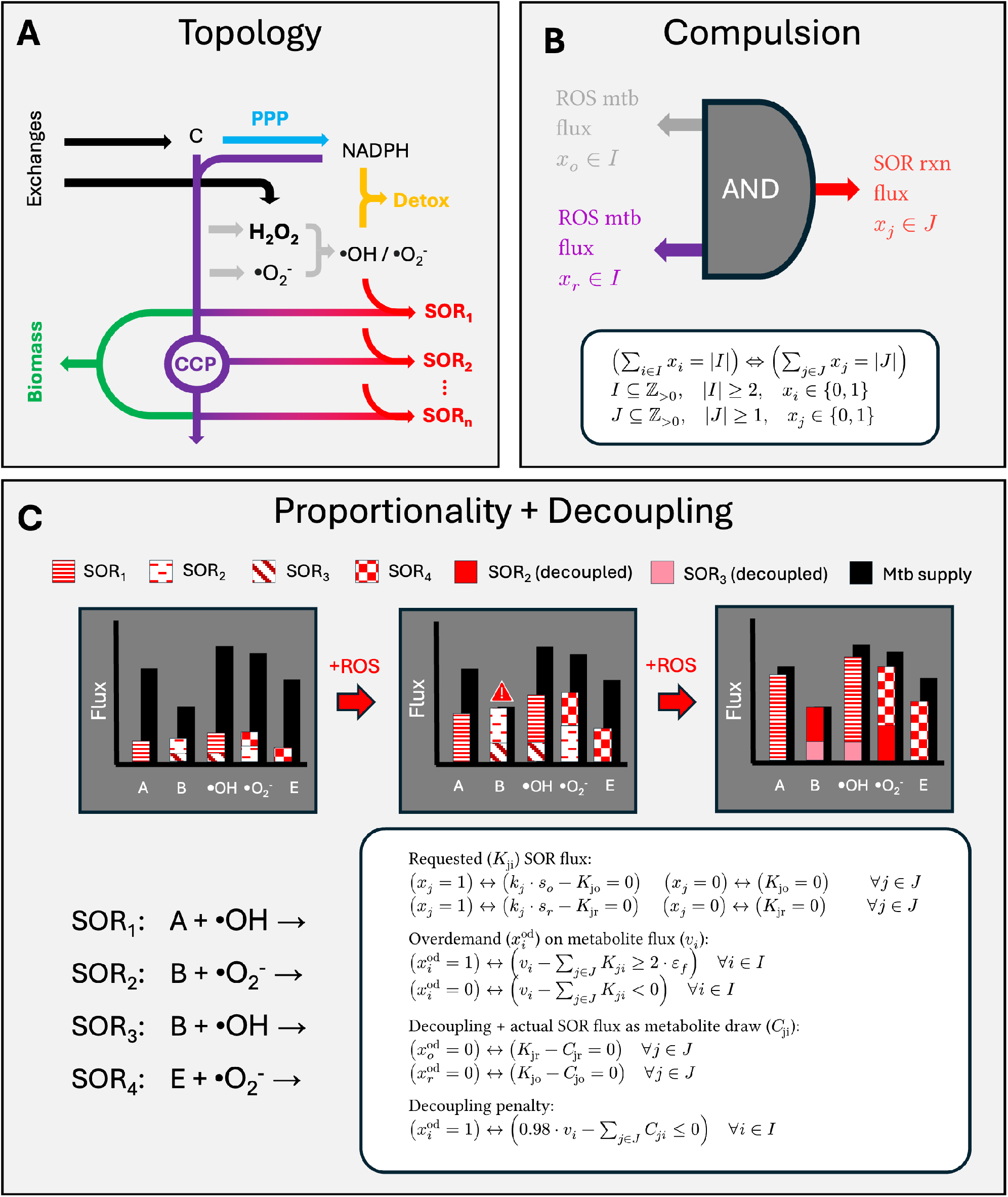
Enforcing kinetic realism in expanded ROS topology via dynamic flux-proportionality constraints. **(A)** Expanded ROS network: spontaneous oxidation reactions (SORs), detoxification, and H_2_O_2_/ROS-generating redox leakage (*topology*). **(B)** AND-gate constraints enforce positive SOR flux only when both reactants have non-zero flux (*compulsion*). **(C)** Intended behavior of rate-constrained reactions (SORs, detox) via reactant draw, illustrated with SORs. First diagram: under rising oxidative stress, SOR flux scales in lockstep, proportional to their rate constants (*proportionality*). Black bars denote maximal supply of metabolites A, B, E, •OH, •O_2_-for a given flux strategy; red boxes show their consumption by four SORs. Second diagram: further stress hits the supply-demand limit of metabolite B by SOR2/3 (red warning triangle), permitting decoupling from lockstep (*decoupling*). Third diagram: additional stress may then be managed by alternative flux distributions or remaining SORs with space capacity, provided SOR2+3 dominate B consumption. Continued increase of oxidative stress eventually becomes infeasible (*death*).

Representative version of the SOR portion of model topology (**Figure S2A**) was simulated to validate **Rules 3-5**, assuming forced or relaxed uptake of ROS (**Figure S2B-E**). The median scaled fluxes of each reaction were in the same proportion as their *k* rates with/without forced uptake, respectively (model with dynamic constraints and decoupling; **Figure S2B-C**). This effect is impossible to randomly achieve based on the non-constrained models (model without dynamic constraints; **Figure S2B-C**). It was also obvious that without **Rule 5** and under conditions of forced ROS uptake (model with dynamic constrains but without decoupling, **Figure S2C**) the model can sample a fraction of the decoupling-capable model’s range. This effect was also highlighted from the absolute sampling ranges of the models (**Figure S2D-E**). Around 45% of samples have perfect reaction lockstep in the decoupling-capable model and only 16% in the decoupling-incapable version when ROS uptake is forced (**Figure S2C**) while that number is 40% for both models without forced uptake (**Figure S2-B**). Overall, the results conclusively validate **Rules 3-5** and simultaneously prove that decoupling improves the absolute sampling range while also improving proportionality.

We implemented our first principles in model BT, yielding the *Base + Topology + Constraints* (BTC) model. A whole genome knockout (KO) screen was performed on all three models in LB medium to test their ability to grow without nutrient limitation while emulating experiments in the absence (Hashimoto et al., 2005) or presence (Roth et al., 2022) of H_2_O_2_ (see Suppl. Data). No correct predictions were exclusive to model BTC. The model had an increased error rate of about 2%, consistent with the penalty coefficient of **Rule 5** granting 2% leeway for practical reasons. GapSplit was then used to sample the three models, also in LB medium but in the absence of H_2_O_2_, by emulating the culture conditions from Hashimoto et al. (2005). The results (**Figure S3**) showed an almost exclusive ability of BTC to generate persisters implying a baseline level of stress but where all models’ samples were survivors – defined as having ATP maintenance flux of 3.15 or above. At the same time BT/BTC models showed a distinct transition in growers at 20% of maximum biomass, prompting us to sub-divide these growers into *bottom 80% growers* and *top 20% growers*. This observation simultaneously validates using the latter to delineate an untreated centroid reference vector for hysteresis enforcement (see Methods).

### Persisters predominate survivors in minimal medium, but growers dominate culture metabolic activity

Emulating exposure to H_2_O_2_ in MOPS medium allowed us to compare how our models’ metabolite turnover fluxes (see Methods) compare against paired metabolomics data, generated by Jozefczuk et al. (2010). This experiment, also corrected for hysteresis due to the short treatment duration, predicted around 20% growers and 80% persisters – all survivors. Since simulated flux and total metabolite abundances are neither in the same units nor scale and because our simulations represent discrete cellular states while the metabolomic abundances are bulk culture measurements, we relied on comparing between pairwise ratio matrices of metabolites (**Figure 3A**). As such, generally low cosine similarity was expected and observed but we were able to see how well growers and persisters individually match metabolomics data (**Figure 3B**). Growers better reflect the data than persisters, and the top 20% of growers best capture metabolomics in both conditions. Furthermore, the diminished difference in cosine similarity between 0 mM and 5.88 mM H_2_O_2_ in growers reflects the metabolic hysteresis of a pre-exposure state dominated by growth, as expected. The Base model, although validating better, showed no difference in how well growers and persisters explain the data (**Figure 3C**), implying a mechanistic indifference to stress.

**FIGURE 3.**
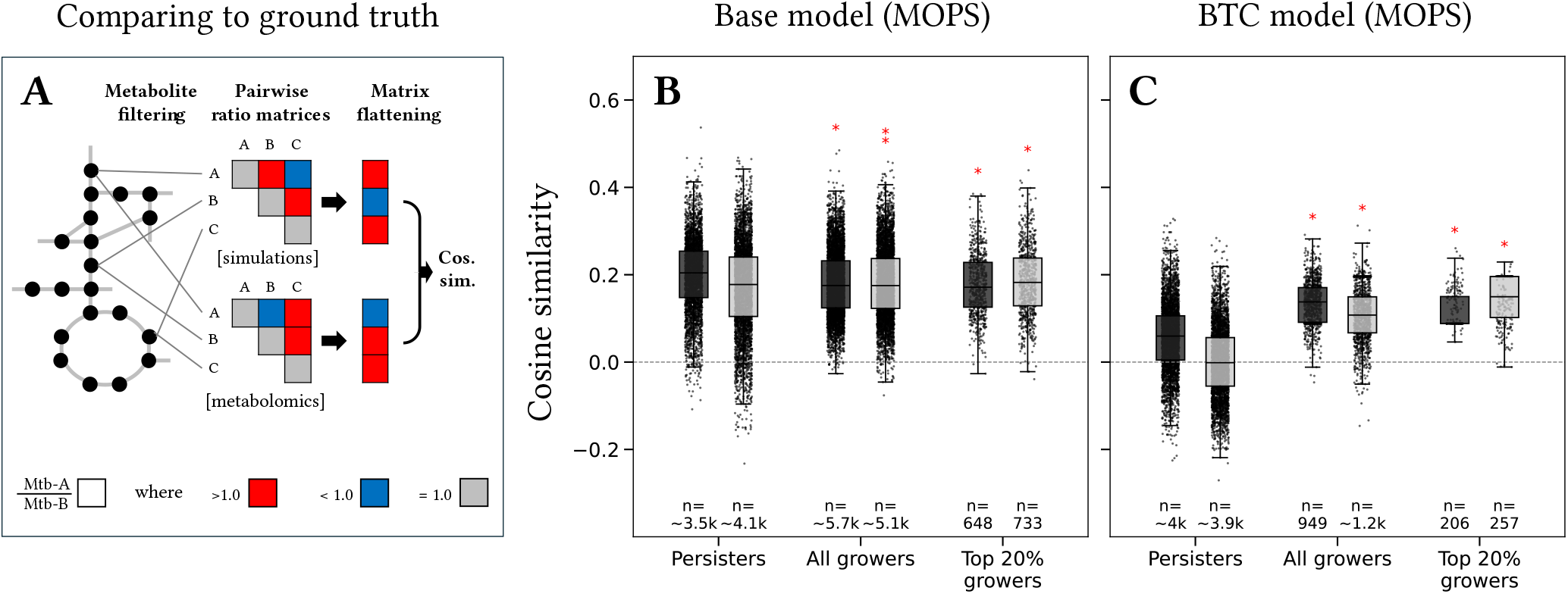
Model BTC persisters and growers show distinct cosine similarity modes against metabolomics data under oxidation stress. Metabolomics data from Jozefczuk et al. (2010), processed per our Methods, were compared against Base/BTC models in MOPS medium sampled with GapSplit, emulating the study conditions (0 and 5.88 mM H_2_O_2_). Simulations were subdivided into persisters, growers, and Top 20% of growers (see Figure S3). **(A)** Method overview: metabolites are filtered per the Methods and intersected with those measured by Jozefczuk et al. (2010) pairwise ratio matrices of fluxes and the measured abundances are then computed and compared by cosine similarity. **(B, C)** Cosine similarity against ground-truth data for Base and BTC, respectively. Significance by t-test with Bonferroni correction: * p<0.05, ** p<0.01.

The Base model results indicated limited capacity to differentiate between metabolic states representing optimum biomass production versus all others. The higher cosine similarity appears to stem from indiscriminate and blind sampling of the entire solution space, which inevitably finds more well-validating solutions. The BTC model, although validating slightly worse in all numeric validations thus far, showed the highest propensity for discrete sub-populations and sensitivity to oxidative stress while its constraints passed all quality controls. As such it was technically a success and could be probed for ROS responses in more depth.

### A loss of persisters causes the bimodal killing pattern in *E. coli*

Evaluating the response of model BTC to a gradient of H_2_O_2_ exposure in growth-limited medium was done as before (Figure S1) to emulate the M9 and H_2_O_2_ conditions from (Brandi et al., 1987), whose data was adapted in **Figure 4A**. Model BTC formed an undeniable bimodal killing pattern in response to oxidative stress (**Figure 4B**). The pattern is qualitatively different from Brandi et al. (1987)‘s but shows the same features – initial decline, a recovery phase, a subsequent decline, and a final small recovery which was outside our scope. Mode 1 killing here corresponds to 0 and 0.23 mM H_2_O_2_. The recovery phase corresponds to 0.47 and 0.94 mM. Mode 2 killing corresponds to: 2.18 to 9.99 mM. Our earlier findings showed us that, at least when it comes to simulations, Mode 1 killing and the Recovery phase are two sides of the same coin (Figure 1). We then broadly classify the state of metabolism in the range 0-2 mM H_2_O_2_, as Mode 1 and 2-10 mM as Mode 2, both defined by the extent of spare detox capacity.

**FIGURE 4.**
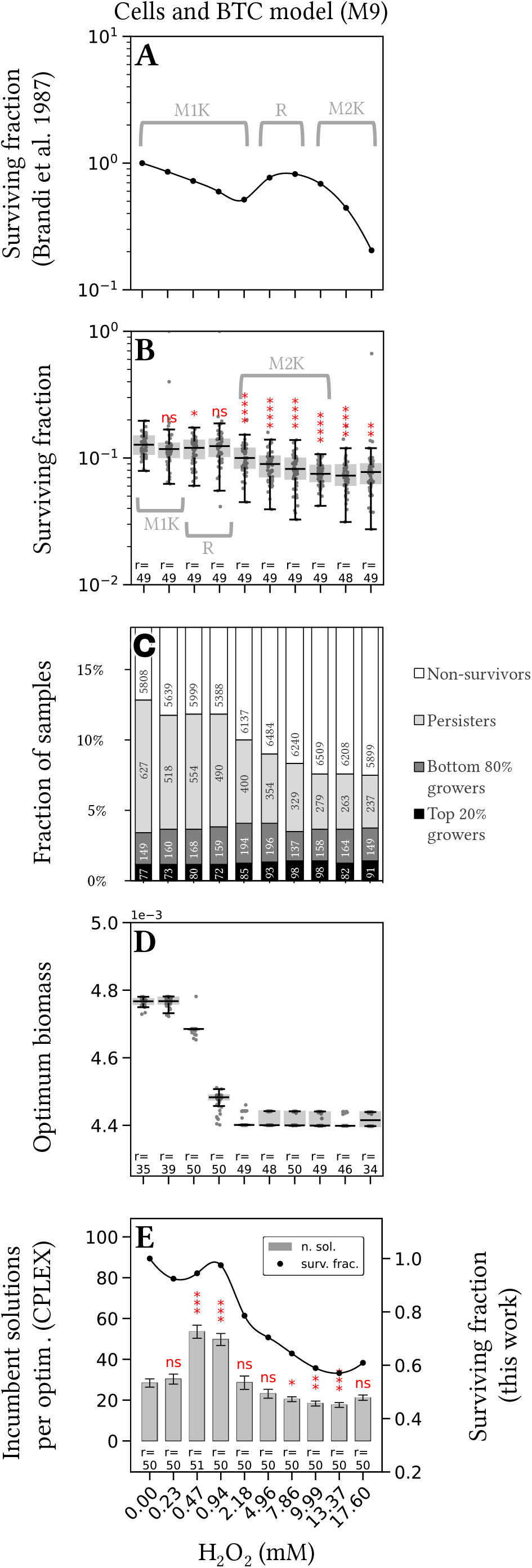
Genomescale models with expanded ROS topology and dynamic constraint system exhibit H_2_O_2_-induced bimodal killing through loss of persisters. The experiment of Brandi et al. (1987) was emulated in M9 medium with the BTC model. **(A)** Digitized raw data (WebPlotDigitizer 4.7), with Mode 1 and Mode 2 killing (M1K, M2K) and the recovery phase (R) – annotated based on the current convention in the field. **(B, C)** GapSplit sampling with hysteresis filtering as in Figure S1. **(B)** Survivor fractions, with killing modes and recovery annotated: Mode 1 (M1), 0-2 mM H_2_O_2_ Mode 2 (M2), 2-10 mM. **(C)** Pooler fractions of non-survivors, persisters, and growers across replicates. **(D, E)** Independent biomass-optimization experiment on the same model using alternative Markov chains through opportunistic parallelism (see Methods). **(D)** Biomass optima; **(E)** mean number of incumbent solutions per optimum from CPLEX logs, with median survivor-sampling results overlaing on a secondary (linear-scale) y-axis. Statistics: panel B, t-test (uncorrected, small n); panel D, Mann-Whitney with Bonferroni correction. ns p≥0.05, * p<0.05, ** p<0.01, *** p<0.001, **** p<0.0001.

Plotting the sub-populations as proportions of all samples showed that the increase of non-survivors came mostly at the expense of persisters (**Figure 4C**), while the numbers of growers (especially the top 20%) is mostly invariable. This invariability is best explained by growth, unlike persistence, being more nutrient-as opposed to oxidation-limited. The consistency in the proportion of growers means that they must be employing strategies mostly sufficient at dealing with oxidative stress. Our work assumes that cells begin treatment as growers, which then stochastically diverge into alternative metabolic decisions constrained by steady state. Persisters, we reason, represent incumbent growers which were unable to maintain steady state (and ATP maintenance) without being simplified by the laws of thermodynamics or the solver. Because of this network failsafe property, persisters tell us little about oxidative stress mechanistically without us being able to observe the transition directly which is difficult without dynamic FBA. Fortunately, persisters due to some heterogeneity, can be considered incumbent solutions, alongside growers, as part of the model’s decision tree. Therefore, if growers do not use a one-size-fits-all response to all treatment intensities, we could use them to identify the decision tree branches (and its failure modes) of all samples. To start off, we performed biomass-optimizations across the gradient of H_2_O_2_ concentrations using CPLEX’s opportunistic parallelism, generating replicates that represent independent optimization paths. The optimum values of the other two models (B and BT) were insensitive to H_2_O_2_ (data not shown). The biomass optima of model BTC were clearly sensitive to oxidative stress despite of the small magnitude drop (**Figure 4D**). The corresponding number of incumbent solutions doubles at the concentrations where sampling formed the recovery phase (**Figure 4E**), supporting the previously theorized threshold-driven regime switching between Mode 1 killing and Mode 1 recovery.

### Persistence is shaped by Ahp, growth is supported by MsrA/B

We anticipated the drop in persisters being due to detoxification failure. Principal Component Analysis (PCA) on detoxification fluxes identified the importance of MsrA/B and Ahp to the respective PCs (**Figure 5A**). Only 9% of persisters used a combination of the two MetSO reductases, while that number was 99.8% in growers. As seen from the PCA, MsrA/B activity is then important to growers but is more moderate in the top 20% among them, implying a somewhat different strategy there. Ahp only defined broad modalities in all survivors with some stratification in persisters. We traced the changes in scaled detox usage of these reactions as a function of H_2_O_2_ in survivors (**Figure S4**). It was expected that persisters and survivors are similar, due to the dominance of the former in the latter. **Figure S4A-B** shows Ahp cleanly corresponding to Modes 1 and 2 for these sub-groups, while being less important to the bottom 80% of growers (**Figure S4C**). Interestingly, the top 20% of growers heavily relied on Ahp (**Figure S4D**). MsrA activity, which is virtually indistinguishable from MsrB (data not shown), generally went up with H_2_O_2_ in survivors excluding persisters (**Figure S4E-F**). As expected the bottom 80% of growers relied extensively on MsrA (**Figure S4G**) while the top 20% used MsrA more moderately (**Figure S4H**). Perhaps the most curious result was the top 20% of growers, whose numbers were mostly invariable, closely resemble persisters in terms of Ahp preference. It thus seems likely that the best and safest fitness strategy is to prevent ROS formation, explaining the use of Ahp which targets H_2_O_2_ directly before it can yield more dangerous radicals in both persisters and top 20% of growers, especially in Mode 1. By contrast, MsrA/B must have more of a damage-control role in most growers, especially in Mode 2, particularly due to its MetSO substrate being downstream of the oxidation damage. The results are so far consistent with Mode 1 being associated with damage prevention, while Mode 2 is focused on damage control.

**FIGURE 5.**
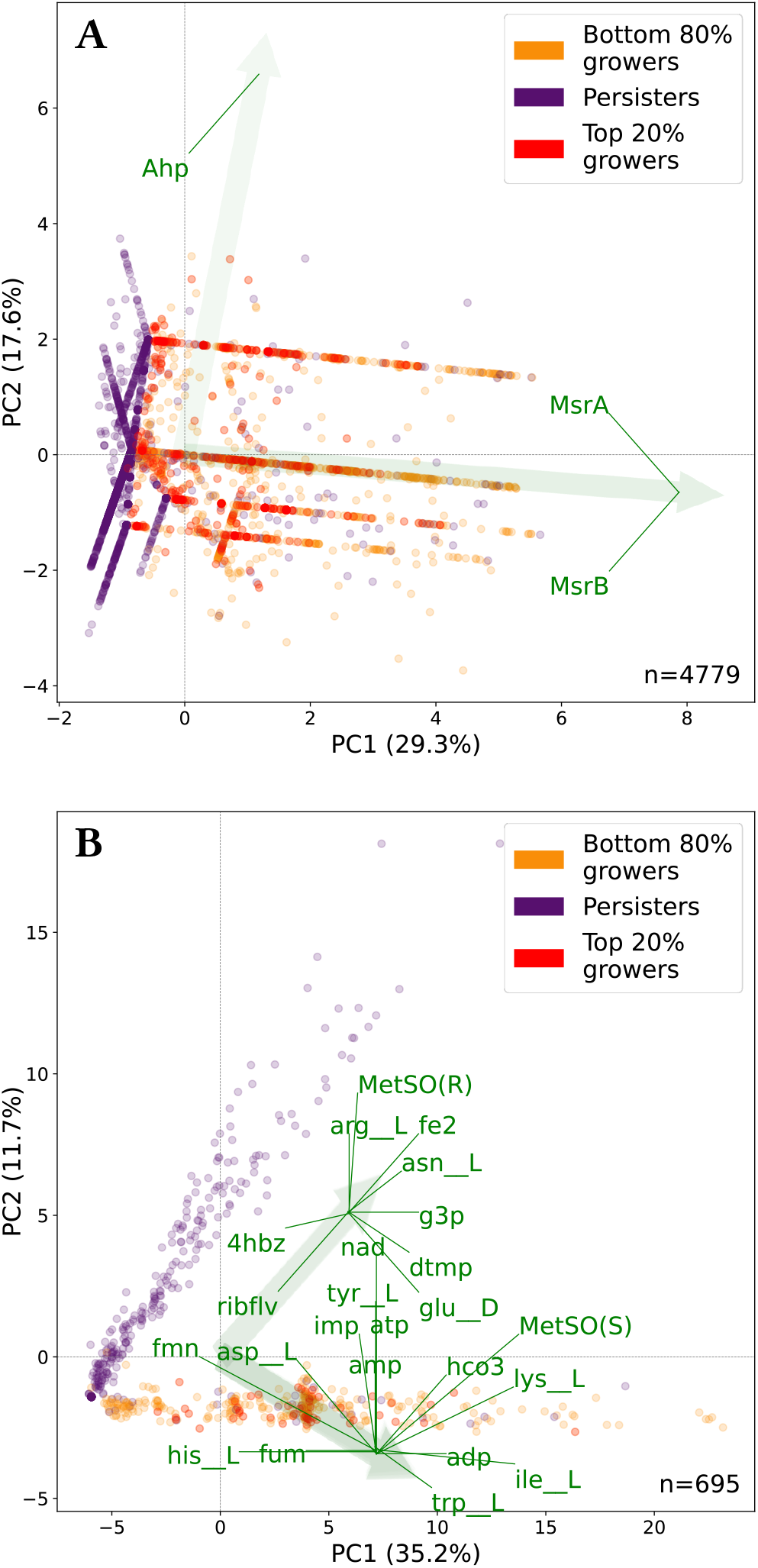
Distinct detoxification (MsrA/B + Ahp) and SOR activity patterns in growers versus persisters. Hysteresis-corrected sampling flux vectors from BTC (M9) were filtered to detox **(A)** or SOR **(B)** reactions and analyzed by PCA. Variables were clipped to the 5th-95th quantiles per variable (outlier removal), so sample counts (n, lower right) degenerate with number of variables. PCA used z-score scaling (mean-centered, divided by standard deviation). Loading vectors (green arrows, labels) contributing ≥ 65% (panel A) or ≥ 95% (panel B) to PC1/PC2. Arrows are length-scaled for visibility and not to absolute loading magnitude. The biomass values for the plotted samples were used for coloring according to persisters (purple), bottom 80% growers (orange), top 20% growers (red) (see Figure S3). All colors and points on the plot have the same opacity of 20% to allow see-through.

We also expected persisters to have failed to distribute ROS along safer SORs. PCA on the SOR usage (**Figure 5B**) revealed both persisters and growers sharing in the spontaneous oxidation of Fe^2+^, G3P, 4-hydrobenzoate, DTMP, riboflavin, and the following amino acids: Arg, Asn, D-glu, Met (as methionine-R-sulfoxide). Growers additionally exhibit oxidation of AMP, IMP, HCO_3_, ADP, ATP, FMN, NAD and the following amino acids: His, Ile, Trp, Tyr, Lys, Met (as methionine-S-sulfoxide). These extra oxidations in growers may indicate that the respective metabolites are rendered more available to SORs due to growth requirements, so we used the same approach to map SOR preference in growers on a per-concentration basis (summarized in **Figure S5**). The most oxidized amino acids across conditions are Lys, Ile, and Trp, followed by His and then Met (as S-sulfoxide). Exclusive oxidations to Mode 1 growers are: riboflavin, ATP, orotidine 5’-phosphate, glycerol 3-phosphate and fructose 6-phosphate. The spontaneous oxidation of several metabolites is exclusive to Mode 2 growers: Arginine, acetate, CMP, cobalt, (R)-Propane-1,2-diol, FeS clusters, reduced riboflavin, D-ribose. As such, the oxidation activity of these metabolites is likely more impacting on fitness than that of their Mode 1 counterparts.

Taken together, these results lead us to hypothesize that Ahp is preferentially used for damage prevention and as such is important to persisters. On the other hand, most growers may be better at picking more favorable SOR usage rather than avoiding damage which is what only the top 20% of growers do. All growers, by definition rely on some MsrA/B to regenerate methionine, whose oxidation is likely but also a direct competitor of biomass. To confirm, we filtered the solution space of our model based on detox reaction usage, essentially simulating a detox KO study in the absence of catalases (**Figure 6**). Testing the detox reactions for their individual effects on survival (**Figure 6A-D**) revealed that only Ahp could single-handedly rescue survival in Mode 1 (**Figure 6C**), although it was clear that the combination of thioredoxin reductase (Trx-HPR) + glutathione peroxidase (GPx) + superoxide dismutase (SOD) + spontaneous dismutation (SOD) caused a general monotonic survival baseline (**Figure 6D**) akin to a model exhibiting only Mode 2 metabolism. Filtering out detox reactions one at a time (**Figure 6E-H**) revealed that Ahp is what causes the survival recovery phase (**Figure 6G**). It seems like Mode 1 is the addition of a recovery phase on top of the otherwise monotonic baseline of Trx-HPR/GPx/SOD/SD. There seems to be a potential combination of detox reactions that is required to support growth (**Figure 6I-P**) but it was clear that Ahp is not important to growth (**Figure 6O**).

**FIGURE 6.**
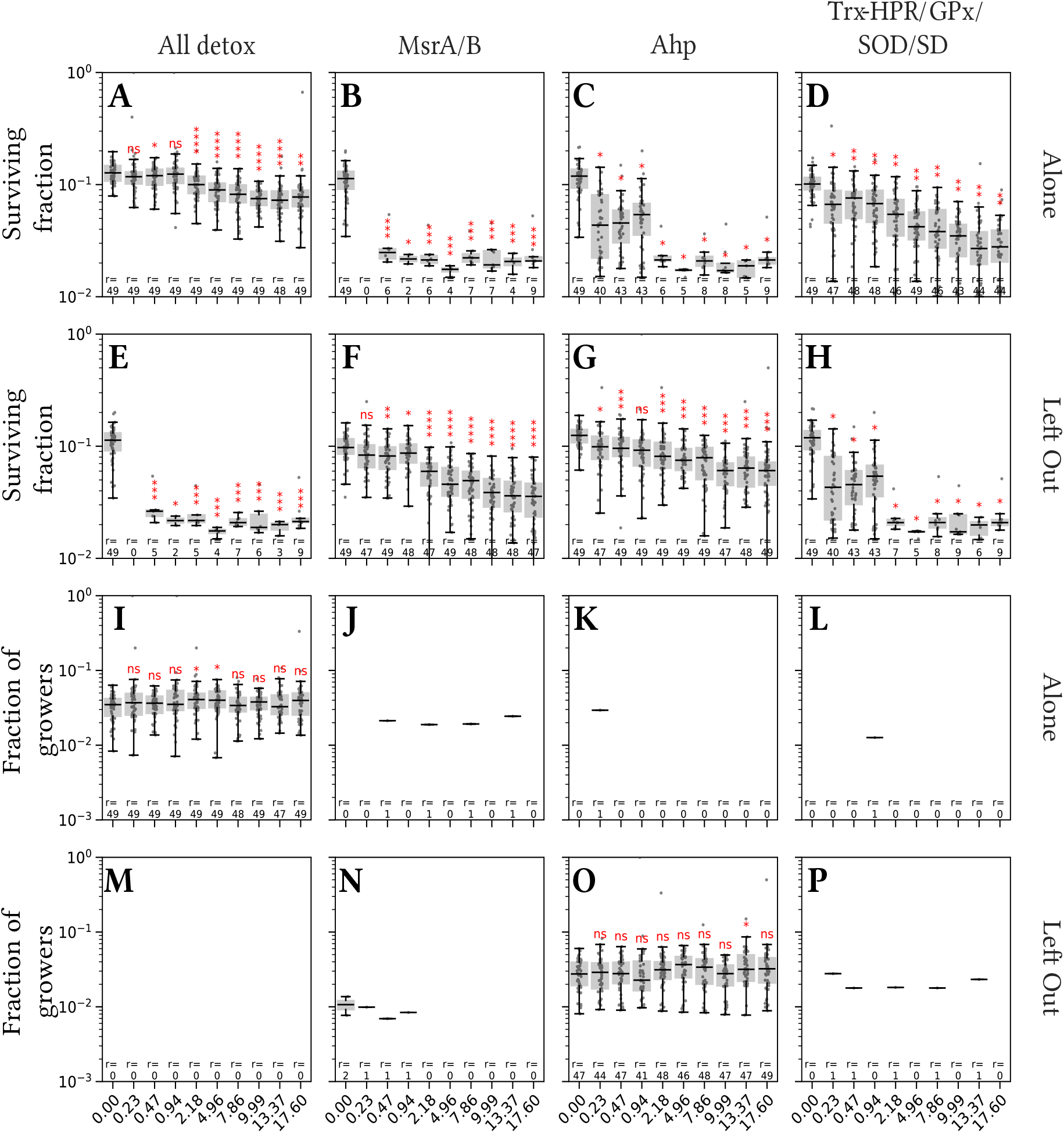
Ahp drives the M1R phase and alone ensures survival up to 2 mM H_2_O_2_. BTC (M9) samples were filtered by the hysteresis vector distance-based cutoff criterion (see Methods), grouped into survivors and growers, and counted as a frequency of all samples. Groups were further filtered by detox system: all reactions, Ahp, MsrA/B, or Trx-HPR + GPx + SOD + spontaneous dismutation (SD). **(A-D)** Survival when each system is solely available. **(E-H)** Survival when each is the only one disabled. **(I-L)** Grower fraction when each system is solely used. **(M-P)** Grower fraction when each system is exclusively used. Significance by t-test (uncorrected): ns p≥0.05, * p<0.05, ** p<0.01, *** p<0.001, **** p<0.0001.

We further dissected the Trx-HPR/GPx/SOD/SD detox systems which were already found to ensure a monotonic drop in survival (**Figure S6**). It was found that GPx and SD could single-handedly ensure survival at low and high H_2_O_2_, respectively (**Figure S5A-H**) and none of the shown combinations could ensure growth per se.

Our findings can be summarized as follows: the most optimum fitness strategy is to prevent ROS formation through Ahp usage. This is less likely in growers but common in persisters which were growers that have likely failed to route flux through safer SORs, thus granting themselves more options for survival. Persisters dominate and shape the bimodal killing pattern, explained mostly through their use of Ahp. Growers may be able to distribute ROS among safer SORs particularly which is particularly important in Mode 2, but more and more MsrA/B is then needed for damage control due to methionine oxidation being critical. Msrs do not single-handedly explain growth but seem to be most important for it, in association with another detox reaction.

### Gluconate shunt, Entner Doudoroff pathway, and higher productivity aide Mode 2 fitness

Next, we investigated broader metabolic changes, associated with the two modes to distinguish the low H_2_O_2_ associated with Mode 1 (0-2 mM H_2_O_2_, low) versus the high H_2_O_2_ associated Mode 2 (2-10 mM H_2_O_2_, high). We compared the metabolite turnover fluxes (see Methods) between the two treatment ranges. We used survivors, persisters, growers and the top 20% of growers between the low and high H_2_O_2_ phases (**Figure 7**).

**FIGURE 7.**
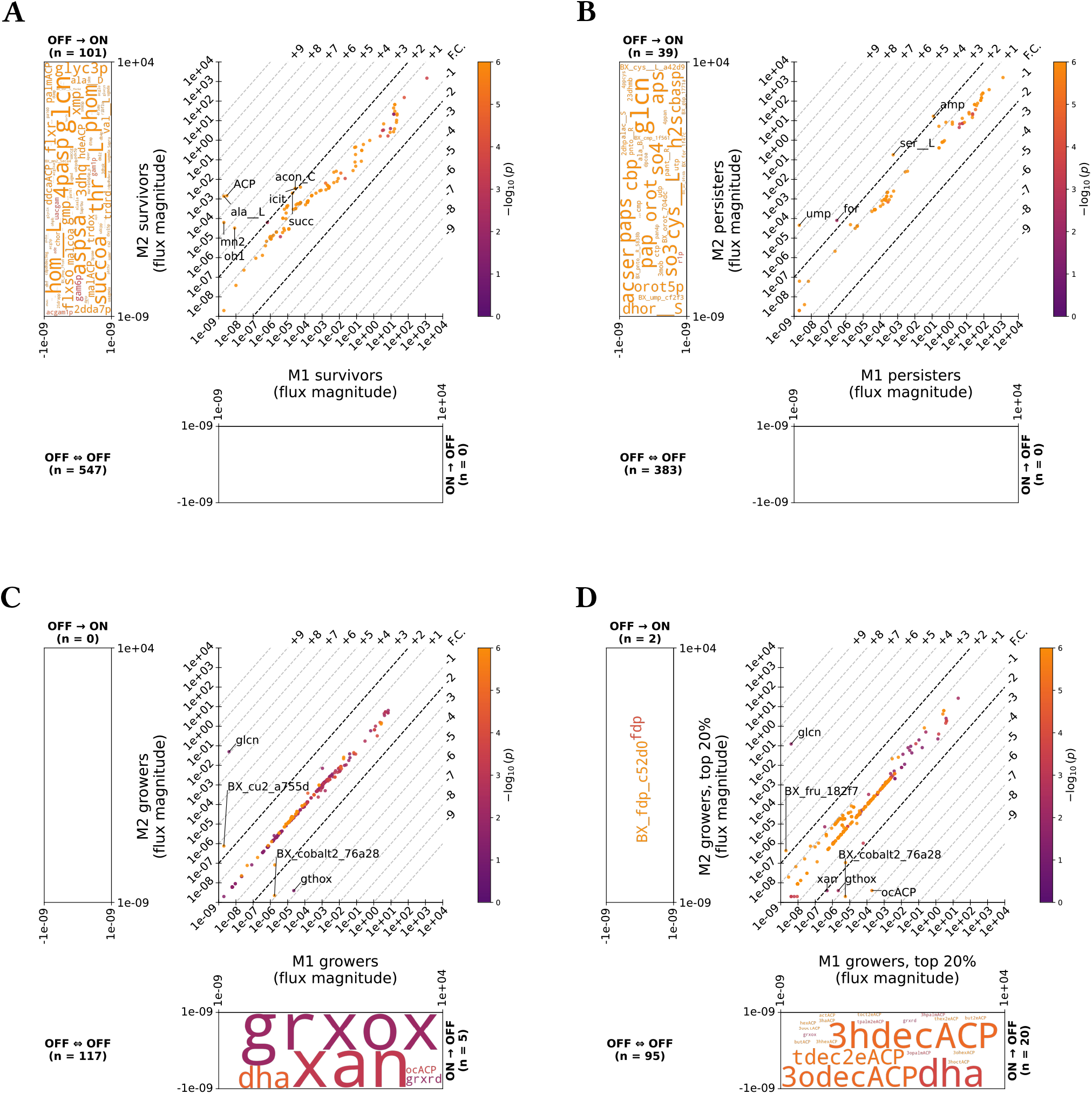
Handling Mode 2 oxidative stress raises metabolic productivity across all survivors; growers additionally engage the gluconate shunt. Flux vectors from BTC (M9) were filtered by the hysteresis vector distance-based cutoff criterion (Methods); metabolite turnover was computed for survivors **(A)**, persisters **(B)** all growers **(C)**, and top 20% of growers by biomass **(D**, see Figure S3). Samples were split by H_2_O_2_ range before comparison: Mode 1 (M1, x-axis) is 0, 0.23, 0.47, 0.94, or [0, 2 mM); Mode 2 (M2, y-axis) is 2.18, 4.96, 7.86, 9.99, or [2, 10 mM). Flux magnitudes are expressed in mmol/gDW/h. Plot areas are colored by p-value (color bar); only metabolites passing the Mann-Whitney significance threshold (p ≤ 0.05) are shown. Cytoplasmic metabolites were used, except those lacking a cytoplasmic version. The scatter shows per-metabolite magnitude change between modes; the center diagonal marks no change, thick dashed lines mark changes within two orders of magnitude (metabolites beyond are labeled). Wordclouds show metabolites transitioning ON →OFF (bottom) and OFF→ON (left) from M1 to M2, where ON/OFF is defined by a single flux value higher than the feasibility tolerance (1 × 10^−9^); word size reflects relative ON-state flux. The lower-left corner shows metabolites OFF in both modes.

Considering all survivors (**Figure 7A**), 101 metabolites are used exclusively in high H_2_O_2_ and exhibit several orders of magnitude higher turnover flux than metabolites shared in low H_2_O_2_. Among these metabolites is manganese, which is oxidized by ROS but is otherwise only used directly in the biomass function. Interestingly, this metabolite effect is only statistically significant in general survivors and not in any of the other sub-populations. A similar effect was evident for L-alanine, isocitrate, cis-aconitate, hydroxide, and acyl carrier protein.

Persisters (**Figure 7B**) used 39 metabolites exclusively in high H_2_O_2_ and had slightly elevated turnover of UMP, formate, L-serine, and AMP. The result of having more metabolites used in the higher H_2_O_2_ range, both in survivors and persisters, is consistent with the Mode 2 being relatively more focused on production, repair, and replacement. It also explains that the majority of NADH must be going to repair or even ATP synthesis since it is not used by Ahp (**Figure S4A-C**) nor biomass.

All growers (**Figure 7C**) varied less with H_2_O_2_. Growers in high H_2_O_2_ use five fewer metabolites, have sever orders of magnitude more D-gluconate turnover, somewhat elevated spontaneous Cu^2+^ oxidation, decreased spontaneous Co^2+^ oxidation and lower turnover of oxidized glutathione.

The top 20% of growers (**Figure 7D**) were mostly akin to all growers. The top growers of the high H_2_O_2_ range make exclusive use of D-fructose-1,6-phosphate, and its associated SOR. D-gluconate also has severn-eight orders of magnitude more turnover in the high treatments. Instead of increased Cu^2+^ SOR activity, the top 20% of growers in the high treatment have an increased spontaneous D-fructose oxidation. Similarly to all growers, a decreased oxidized glutathione turnover, lower spontaneous oxidation of Co^2+^, lower xanthine turnover (which is unused by all growers in Mode 2, statistically). Uniquely, the top 20% of growers in the higher range had around four orders of magnitude less acyl carrier protein-bound octanoyl turnover. Mode 2 in this subpopulation was also more aggressive in completely avoiding metabolites, with 20 metabolite turnovers being completely switched off as opposed to the five in all growers. These metabolites are mostly a superset of the corresponding group in all growers, excluding xanthine and acyl carrier protein-bound octanoyl. The extra unused metabolites were exclusively bound to this protein. Taken together, these results mean that the top 20% of growers are a highly specialized versions of the general growth strategies. These high performers seem to avoid more of their fatty acid synthesis pathways, represented by acyl carrier proteins, in Mode 2 and at the same make use of fructose. It seems that all growers rely on a significant increase of D-gluconate turnover, decreased spontaneous oxidation of cobalt, and decreased oxidized glutathione turnover.

Comparison between persisters and all growers, unsurprisingly, revealed more than 400 extra metabolites being used in the latter for both modes (**Figure S7A-B**). Growers, by definition, are high producers. As such, it was more useful to look at what was exclusive or unique to persisters. For example, persisters use less D-ribulose 5-phosphate and D-xylulose-5-phosphate than growers in Mode 2, showcasing a decreased reliance on the extended Pentose Phosphate Pathway (PPP) which is not surprising, considering their smaller need for NADPH for biomass and Msrs. Persisters, unlike growers, make use of 6-phospho-D-glucono-1,5-lactone, O-phospho-L-serine, and 3-phosphohydroxypyruvate. The 3-phosphohydroxypyruvate is a byproduct of the regeneration of NADH by 3-phospho-D-glycerate, a glycolysis metabolite, via phosphoglycerate dehydrogenase. The produced 3-phosphohydroxypyruvate is then converted to O-phospho-L-serine through the phosphoserine transaminase and later to L-serine through phosphoserine phosphatase. This short pathway, exclusive to persisters, therefore represents a direct way to produce NADH directly from glycolysis for multiple applications including powering Ahp.

The differences between the top 20% and bottom 80% of growers in Mode 1 consists of the top 20% using 39 fewer metabolites, in addition to avoiding the spontaneous oxidation of reduced glutathione and its SORs, while also using less propionate and propanoyl-CoA in the lower range (**Figure S7C**). The results are similar in the higher range where the top 20% growers use only 15 more metabolites than bottom 80% of growers, consistent with the effects of more inhibited production. The only other difference between these groups of samples is the top 20% of growers having eight orders of magnitude less 2-dehydro-3-deoxy-D-gluconate in Mode 2. This metabolite is part of the Entner Doudoroff (ED) pathway, which can shorten PPP to feed back into glycolysis, thus generating less NADPH. This result could explain why the best growers mostly avoid MsrA/B usage and instead rely on Ahp. There are roughly twice as many bottom 80% as there are top 20% of growers. This implies that Msr usage is the more primed ROS detoxification mode in culture before stress exposure.

## Discussion

Our approach was unorthodox in that we pre-compiled a list of all reasonably possible solutions for each condition independently, and treated all of them as related within set Euclidian distances. We are thus accounting for both bet-hedging and hysteresis, where individual samples are more akin to strategies as opposed to separate cells. The implication of such an approach is that independent strategies could be mapped onto decision tree branches based on sets of their characteristics. This reveals stochastic likelihoods of making different metabolic decisions as an emergent property of the network and its constraints. Perhaps even more important is the consequence of this that both growers and persisters, which are survivors that do not invest in growth (Radzikowski et al., 2016), can be treated as incumbents on the branches of this theoretical decision tree. This perspective allows us to use growers to identify the sets of characteristics specific to each tree branch where this would not be possible from persisters alone. Persisters are also characterized by heterogeneity but are mostly single-minded in terms of function, focusing mostly on covering ATP maintenance. We could then consider that for each persister’s position in the solution space there was initially a failed grower that could not satisfy its constraints given its unique stochastic flux choices and was thus resolved into a simpler solution – a persister. The cause of persistence in living cells according to Radzikowski et al. (2017) is a collapse in metabolic fluxes. After all, we do expect the samples to start off as growers before treatment as growth is what unstressed cells are seemingly optimizing for (Goelzer et al., 2015; Harcombe et al., 2014; Molenaar et al., 2009; Schuetz et al., 2012). Likewise, we expect that for each dead sample there was initially a failed persister. This allowed us to use growers as solution space representatives and offered us a glimpse of the network-wide changes during oxidative stress, where simpler solutions (dead cells, and to a lesser extent persisters) would have otherwise been mechanistically uninterpretable. Despite of the proportion of growers being remarkably invariable in our survival experiments, we found them to offer good explanation of the drop in persisters which our work is focused on. Therefore, we assert that pre-existing bacterial heterogeneity of high-performing growers biases the formation of persisters under stress. Evidence exists against the alternative *slow growth hypothesis* (Goormaghtigh & Van Melderen, 2019) which, along with our metabolomics validation, justifies our usage of only high-performing growers in the untreated conditions which also biases hysteresis. An alternative interpretation of our metabolomics validation (**Figure 3**) is that growers may be rare but dominate the culture’s metabolic activity. Another benefit of our strategy on top of the general optimization approach was that we started with the entire solution space in the form of GapSplit samples. We simply filtered solutions for specific characteristics, essentially standing in for KO and even rescue experiments, which was further beneficial for mechanistic interpretation.

The method presented in this work is most likely unsuitable for general use for several reasons. First, running the optimizations was very computationally expensive and took several months even with parallelization onto HPCs. Second, the large models require handling through a special pipeline for loading and running in order to ensure the correct solver and cluster settings are applied when the model is loaded, which our constraint systems are extremely sensitive to. In addition to this, the pipeline handles gene knock outs as they work slightly differently from normal COBRA models. Third, which became evident from the metabolomics and whole genome KO validations, our models come with a relatively small but noticeable trade off in numeric validation, undoubtedly a function of focusing on a subset of iJO1366′s solution space. A larger and more uniform solution space is bound to produce more samples with higher accuracy, since most of the model’s topology has nothing to do with ROS, even if that higher accuracy is blind to the intricacies of ROS. Proof of this was the observation that the solution space of our survival experiments does not approximate actual culture behavior under oxidative stress even with the ROS topology while in the absence of our constraints (**Figure S1**). After all, the general behavior of the culture is its highest emergent property and we specifically designed our five rules of oxidation realism to be as general as possible to avoid fitting data. Implementing the reaction rate constants is unavoidable parameterization in anything but a molecular dynamics simulation since our models have no spatiotemporal dimensions. It is also worth mentioning that, strictly speaking, the reaction rate depends on the *k* rate constant in addition to the concentrations of two reactants [*A*] and [*B*] as a second order rate *k*[*A*][*B*]. Since it is (currently) impractical to calculate the the steady-state metabolite concentrations in GEMs, our SOR and detox fluxes are only approximations. Calculating steady state concentrations from fluxes is difficult without some sort of parameterization of the metabolite half-life *in vivo*, a problem also for the metabolomics validation. Fourth, the choice of ROS topology to reconstruct reflected what can be modeled in iJO1366 and the subset of SORs (and their constraints) must be adapted to the host model.

Large models have more avenues to cheat. Despite of the drawbacks listed above our work proved that the five rules of oxidation realism, in addition to a few minor considerations like H_2_O_2_ dosing, were sufficient to reproduce the general features of oxidation-stressed cultures while offering us insight into the mechanisms and forces at play on metabolism under such conditions. Putting the listed considerations aside, our work shows that there is a metabolic component to the bimodal killing pattern which was previously overlooked in favor of biopolymers (Linley et al., 2012). More so, our results lead us to hypothesize that it is entirely possible for the bimodal killing pattern to initially be caused by metabolism.

We expect real cell metabolism to strive towards steady-state, just like our simulations, as this is the network state with least amount of free energy (Wilson & Matschinsky, 2021). Having this assumption in mind, we offer an explanation of why our modeling can qualitatively recapitulate the bimodal killing pattern and predict a majority of persisters among survivors purely from a model of steady-state fluxes. Both our small theoretical model (**Figure 1**) as well as the BTC model successfully recapitulated a bimodal killing pattern as an emergent network property, also proving that the constraints are only needed insofar as to prevent larger models from cheating. The real driving force behind the metabolic component of the bimodal killing pattern seems to be entirely stochastic and topological, as seen through our small theoretical model, and we suspect the principal driving force is the adherence to steady-state. Looking at the small theoretical model in **Figure 1A** we note that the network is entirely non-deterministic, subject only to *S* · *v* = 0 and variable bounds. This means that the pull of precursors (i.e. carbon) and output of fitness is expected to be random unless it would otherwise endanger steady state. Starting from a small amount of oxidative stress, we note the IQR variation of detox and SOR usage (**Figure 1G-I**) once again highlighting the non-determinism towards fitness as ROS flux can be handled agnostically through both detox and SORs. The stochastic increase of SORs is what causes the initial Mode 1 killing (**Figure 1E**). At this point, the extra pull of precursors to deal with ROS is small and well within the capacity of both central- and alternative-carbon pathways, meaning that the aggregated stochastic fluxes are split 50:50 among the two pathways regardless of bounds. As ROS increases further the capacity limit of central carbon metabolism is inevitably reached and relatively more precursors are bound to go to alternative carbon pathways instead (**Figure 1K**). What causes the sudden bias towards detoxification then (**Figure 1I**)? The key is that alternative carbon pathways can only send an amount of flux back into central carbon pathways equal to the initial flux or it would violate the steady-state assumption. This means that upon reaching the central carbon pathway limit, a bias is created for the alternative carbon branch (detox) in order to preserve *S* · *v* = 0 and handle ROS. This bias for increasing detox is what causes the Recovery phase and is entirely due to balancing fluxes as opposed to trying to improve fitness, which would otherwise be a deterministic behavior. Proof of this explanation is the fact that a bimodal killing pattern is not seen when alternative carbon metabolism has a higher capacity than central carbon’s (**Figure 1B-E**). Alternative carbon metabolism is partially representative of the PPP and an excess of PPP capacity is a characteristic of some cells and has been demonstrated before (Christodoulou et al., 2018). Another parallel to the literature is the effect that at 4 °C there is no Mode 1 killing (Brandi et al., 1987) and likewise when metabolism is inactive (Imlay & Linn, 1986), both of which are consistent with our explanation of it having an enzymatic cause. The Recovery phase ends when the detox capacity has been saturated. At that point the only way to handle the increase in ROS is to keep detox high and simply let more flux go into the SORs, causing the Mode 2 killing, and a bias for SORs. This could also explain the dependence of Mode 2 killing on oxygen in cells seen by Brandi et al. (1987). We thus anticipate that relatively more potential Mode 2 killing solutions would be lost due to being infeasible with respect to *S* · *v* = 0. The real-life equivalent of this effect would be the accumulation of ROS as it cannot move through the metabolic network fast enough, which may increase spillage into the biopolymers given its reactivity. Several sources, reviewed in Linley et al. (2012), suspect Mode 2 killing being associated with biopolymer damage – particularly that of lipids. More importantly, our findings are that Mode 2 killing is not necessarily caused by detox but is characterized by detox being insufficient and this mode’s SOR usage looks nearly identical to that of a model without detox capacity (**Figure 1F-G**) which explains the tendency of solutions in Mode 2 to be focused on production rather than prevention in order to compensate for the damage. A natural conclusion to our observations is that Mode 1 is caused by detoxification granting an initial bonus to fitness in low oxidation stress. It could also be considered that all cells exist in Mode 2 by default, with detoxification modifying the survival baseline (**Figure 6D**) with a Recovery phase, thus yielding Mode 1 (**Figure 6A,G**). As such, Mode 1 has an excess of detox capacity and does not require prioritizing it.

Thus, our work presents the metabolic component of bimodal killing pattern as nothing more but a fundamental thermodynamic effect of this ubiquitous network topology, metabolic stochasticity, and the flowthrough of substance.

Our small model predicted the two modes and identified two respective detoxification patterns. The larger BTC model was able to identify the dominance of NADPH- (mostly MsrA/B) in growers, especially Mode 2 growers. Interestingly, the top 20% of growers instead rely on Ahp, which is akin to persisters, and have only a small dependence on MsrA/B. We suspect that this preference has to do with hysteresis from untreated conditions. Both unoxidized Met and NADPH are biomass components so there must be an obvious preference for using the PPP. Redirecting more flux towards the PPP is easily achieved through the gluconate shunt, which is also the case for real eukaryotic cells experiencing oxidative stress (Schittenhelm et al., 2019). The top 20% of growers also use the gluconate shunt as they too need PPP biomass components. However, these adept growers may redirect some of the PPP flux back into central carbon pathways through the ED pathway, which is a hallmark for improved survival of *Pseudomonas putida* KT2440 under oxidative stress (Chavarría et al., 2013). We suspect that this additional modification gives the detoxification portion of the PPP flux to Ahp and is what makes Ahp-dominant growers stochastically rarer. Overall, growth consists of a lot of moving components which are likely to fail under stress and this is why persisters, and Ahp by extension, dominate our simulations and reality. In fact, Ahp is the principal cause for Recovery according to our data (**Figure 6G**), whereas MsrA/B barely affect survival and are instead needed mostly for growth, as hypothesized (**Figure 6I-P**). It is tempting to challenge this conclusion because MsrA/B is required to recover cells on plates under oxidative stress (Atack & Kelly, 2008; Moskovitz et al., 1995) but it is important to remember that recovery on plates is based on growth. We also suspect that Ahp is how a persister-dominated culture, which is the dominant form of survival *in vivo* (Amato et al., 2014), clears the extracellular environment from H_2_O_2_ but most of the NADH goes to ATP maintenance instead of detox (**Figure S4B**). Uhl & Dukan (2016) have shown that cells (persisters inferred) move from Ahp-dominance to catalase dominance as H_2_O_2_ increases. Finally, we suspect that the increased heterogeneity of solutions in Mode 1′s Recovery phase (**Figure S5C**) has to do with the bias towards detox allowing multiple alternative SOR usages, which also explains the invariabiliy of the number of used SORs to ROS in Mode 1 but not Mode 2.

Our findings could be important to other fields too as fundamental biology effects often emerge in distant species. For example, the Crabtree and Warburg effects link yeast and tumors with mitochondrial dysfunction (Brand & Hermfisse, 1997; DeBerardinis & Chandel, 2020; Diaz-Ruiz et al., 2011; Perez-Samper et al., 2018; Shiratori et al., 2019). Mitochondria are directly linked with disease via uncoupling protein 2 (UCP2) expression (Baffy, 2010; Franceschi et al., 2018; Sharma & Sampath, 2019; Theurey & Pizzo, 2018; Vallejo et al., 2021). UCP2 expression reduces the mitochondrial proton gradient causing thermogenesis and slowing down ROS generation (Brand & Esteves, 2005; Nesci & Rubattu, 2024). There is also a strong relationship between methionine sulfoxide reductases (Msrs), mitochondria and disease (Jiang & Moskovitz, 2018). Slowing down ROS generation is critically important to cancer development (Perillo et al., 2020) as hypoxic cells like tumors are especially susceptible to free radical damage (Wang et al., 2019). We see no reason why the bimodal killing effect or the switch to persistence could not be affecting cells and mitochondria in human disease, especially since it has been observed in eukaryotic cells (Kaneko et al., 1994) and is based on a ubiquitous network topology. Interestingly, only the H_2_O_2_-resistant line (Hpr-4) from (Kaneko et al., 1994) exhibits the bimodal killing pattern, further highlighting the importance of detoxification. The Hpr-4 line contains two mutations in p53. These mutations could be influencing TIGAR function, which has a known effect on NADPH levels, as reviewed by (Lee et al., 2014). All things considered, the bimodal killing pattern of *E. coli* could give us more insight for medicine.

## Summary & Conclusions

We mathematically enforced fundamental rules of ROS and simulation realism as an optimization problem, which allowed us to qualitatively recapitulate the famous bimodal killing pattern and prolonged lag phase, dominated by persisters in *E. coli*. Our findings allowed us to distill the bimodal killing pattern to nothing more than a stochastic effect of the network topology, caused by ROS detox under a steady-state assumption. We theorized that the Mode 1 killing is a continuation of Mode 2 killing but with a more and more active detox system (jointly defined as *Mode 1*). Mode 1 is characterized by detoxification having reserve capacity left. By contrast, Mode 2 is where the detox capacity is at its maximum, causing the metabolism to focus more on repair and replacement of damaged metabolites and overall, more anabolic activity. We identified these behaviors as a natural metabolic trajectory of growers exposed to oxidative stress, where failure to stochastically achieve them leads to falling back to persistence, which is the dominant state, or furthermore – death.

Persistence is mostly focused on minimal metabolism focused on survival through ATP generation and complete lack of growth, thus minimizing the ROS attack surface. This metabolic “safe” mode can expand some of its ATP potential to generate NADH for minimal detox activity, specific to Ahp, which may explain another famous effect of H_2_O_2_ in *E. coli* – the prolonged lag phase, followed by exponential growth. Persisters contrast to growers, which are rare in general but predisposed towards a gluconate shunt that may help them drive more flux into the PPP to power MsrA/B. The best growers occur half as likely and redirect flux from the PPP back into central carbon pathways through the ED pathway, focusing more on NADH generation and Ahp use in a persister-like behavior. The overall dominance of persisters and persister-like behavior is most likely predisposed by pre-existing bias towards central carbon metabolism of all cells.

Death on the other hand seems almost entirely due to losing the persisters and we suspect it may fundamentally have to do with the steady-state of metabolism in Mode 2 killing, and ATP homeostasis in Mode 1 killing. At low stress levels the influx of ROS can be handled through a combination of detox systems and SORs resulting in a larger variation of fitness, biased towards a dip in survival (Mode 1 killing) due to SORs competing with ATP maintenance since persisters have less means to repair themselves. From a non-determinism standpoint, we suspect that the subsequent recovery phase is not due to survivorship bias but happens as a natural network property to preserve steady-state where the only way to handle the extra ROS flux through the system is to use all available pathways, resulting in increase of detoxification due to its reserve flux capacity. This extra detoxification bias will naturally reduce the extent of SORs, even though they are not completely lethal at that point, which increases survival. The exhaustion of this detox capacity brings about the start of Mode 2 which is equivalent to a network without detoxification and the spare capacity to drive it. We suspect that Mode 2 killing is itself due to excess ROS reaching the natural limits of steady-state through the network, resulting in infeasible solutions in our simulations or spilling over to increased polymer oxidation in real life.

Our results form a deeper level of understanding about the dynamics of ROS from a non-deterministic network perspective. We see absolutely no reason why the effects reported by us and others should be exclusive to *E. coli* as they are fundamentally based on ubiquitous network topology and stochasticity. As such, our work may be important to fighting pathogen persistence, or even translating to preventing loss and damage of mitochondria in human disease. Future work could address some questions on a deeper level or even investigate the predicted effects in other systems.

## Methods

### Base model constraints

We used iJO1366 (Orth et al., 2011) as the Base model for *E. coli* metabolism upon which the Base + Topology (BT) and Base + Topology + Constraints (BTC) models were developed. Every model we report has the following shared topological constraints for fair comparison. To promote only aerobic growth, a forced minimum O_2_ uptake equal to the solver feasibility tolerance was set. The lower bound on cobalamin (vitamin B_12_) exchange was set to −0.01, a growth requirement in iJO1366. Aside from the lower bound on O_2_ described above, exchange bounds for water, gases, and inorganic trace ions that should not be growth limiting (CO_2_, H_2_O, O_2_, H^+^, Zn^2+^, Cu^2+^, Co^2+^, Mn^2+^, Ni^2+^, and Mo^2+^) were set to [−1000, +1000], effectively unlimited.

The lower bound of ATP maintenance (ATPM) was set to 0 to permit non-surviving solutions. We observed that sampled solutions follow a binary distribution, with ATPM values of 0 or 3.15; therefore, the biomass reaction was coupled to the ATP maintenance flux through a dynamic constraint so that no biomass flux was possible if ATP maintenance was less than 3.15. This constraint converts the all our reported models into a Mixed-Integer Linear Programming (MILP) problem. Finally, all reactions associated with catalases (CAT, HPI, HPII) were disabled, as none of the experiments our models emulated included adaptation to oxidative stress that is typically required to observe catalase expression (Loewen et al., 1985).

### Network reconstruction of ROS chemistry

All metabolic network modifications to our models are listed in the Suppl. Information file, including disabled and added reactions. The most important modifications begin with splitting the influx of oxygen into two parts by the *O2tex_ep* reaction, replacing the *O2tex* reaction so that 1% of the O_2_ is earmarked for ROS production:

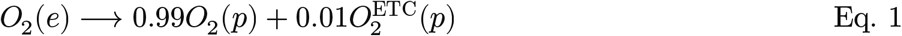

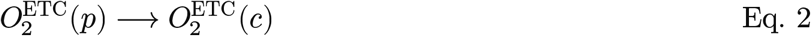

where *ETC* stands primarily for electron transport chain leakage of electrons to produce ROS but also accounts for cytosolic redox reactions contributing to ROS production. We based the percentage of O_2_ to be converted to ROS by this mechanism on mitochondrial research (Chance & Williams, 1956) as this data was not directly available for *E. coli*. According to the authors 1-2% of the oxygen ends up as radicals, prompting us to pick the lower bound as the relationship is dependent on the concentration of oxygen. This cytoplasmic 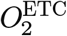 molecule is then used as the oxygen substrate for 14 redox reactions with the effect of producing •O_2_ or H_2_O_2_. Dynamic constraints were used to track the usage of these redox reactions and enforce the proportions of their oxidation leakage counterparts. There are many more known redox enzymes in iJO1366, but actionable leakage data was only found for these 14. The escaped oxygen can also participate in four Fe-S cluster damage reactions, but these aren’t enforced in our model due to the complexity of accounting for specific parameters and optimization requirements not compatible with our approach but treated in a previous study (L. Yang et al., 2019). Most of the ROS-producing reactions we reconstructed produce •O_2_, and some produce H_2_O_2_. These ROS feed into the Fenton reaction and Haber-Weiss cycle, summarized as:

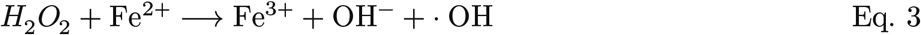

and

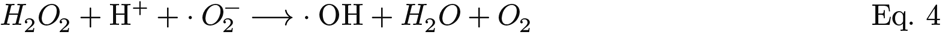

which replace the default iJO1366 reaction FESD1s, which would otherwise convert all H_2_O_2_ to H_2_O.

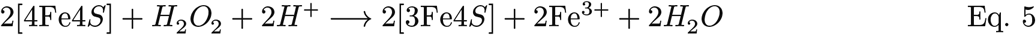

The •OH-generating reactions are the only consuming pathways for H_2_O_2_, leading to ROS stress. Our models’ scope is the metabolome, but a substantial proportion of generated ROS would oxidize macromolecules such as nucleic acids, lipids, and proteins. Because these macromolecules cannot be directly encoded in our model, we instead redirect a proportion of ROS flux representing these reactions into a special demand reaction, one for each radical. We approximate the proportion of ROS that reacts with the small molecule metabolome is equal to the corresponding metabolomic proportion cell dry weight – 3.5% (Neidhardt et al., 1990). The remaining 96.5% of ROS flux leaves through the special demand reactions under control of a proportion-enforcing constraint and is not available to SORs.

We reconstructed SORs based on extensive reviews of spontaneous chemistry for •OH (Buxton et al., 1988) and • O_2_- (Bielski et al., 1985). iJO1366 metabolites were cross-referenced with PubChem, BiGG, MetaNetX and ChEBI. Synonymous entries were merged using the obtained SMILES strings and fuzzy string matching, resulting in a list of synonyms for every metabolite in our model. Optical character recognition (OCR) was used to parse the reaction data from Buxton et al. (1988) and Bielski et al. (1985), and the parsed primary reactants were searched in the list of synonyms to iJO1366 metabolites. We then manually reviewed all potential matches, ensured that all SORs were mass and charge balanced, and filtered out SORs that do not occur under physiological pH or involve dead end iJO1366 metabolites. The remaining curated SORs were added to the Base model to give the topology of the BT and BTC models, and associated rate constants were used to derive respective dynamic constraints. ROS detoxification reactions, also under the control of dynamic constraints, were reconstructed based on manual literature review (Gray & Carmichael, 1992; Lin, 1999; Mishra & Imlay, 2012; Palma et al., 2020; Seaver & Imlay, 2001b; Snezhkina et al., 2019; Tarrago et al., 2012).

### *In silico* culture media

With the exception of exchange constraints described above, we modeled LB, M9, and MOPS media conditions as follows. LB formulation was taken as previously described (Marinos et al., 2020). Exchange bounds were set equal to millimolar concentrations for respective components in all media, resulting in biomass fluxes that represent yield relative to medium composition rather than actual growth rate, as noted in previous work (Marinos et al., 2020). Nevertheless, the maximum biomass flux in our models grown in LB was 2.28/h, close to the measured specific growth rate of 2.1/h in mid-exponential phase corresponding to a doubling time of 20 min (Sezonov et al., 2007).

M9 formulation was based on (2010) by Cold Spring Harbor. The subset of trace elements (e.g. Fe^2+^) that are not provided in excess as described above are based on a separate previous study (Flamholz et al., 2020). We additionally set the bound on 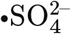 uptake to 1.21 × 10^−3^ mM in M9. MOPS formulation was based on Teknova, CA (product number M2106) as described previously (Jozefczuk et al., 2010) and the same trace elements as accounted for in M9 and any modifications done to the media by the same authors. All *in silico* media formulations and calculations are available in the Suppl. Information file.

### Estimating H_2_O_2_ influx rates

Because H_2_O_2_ is detrimental to culture growth and survival, both in biological reality and our ROS-chemistry expanded models, its uptake must be forced through constraints; otherwise, optimizations of biological objective functions such as biomass will avoid its uptake. H_2_O_2_ uptake rates are based on spontaneous diffusion across the cell membrane. The rate of H_2_O_2_ diffusion across the cell membrane was modeled using ordinary differential equations (ODEs) and parameterized, inspired by the experiment of (Seaver & Imlay, 2001a). This was done by treating a cell culture as two compartments – the combined volume of the cells and media. The extracellular medium volume was calculated as:

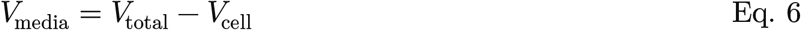

The average influx of H_2_O_2_ is computed by differentiating the equilibration of H_2_O_2_ concentration on both sides of the membranes over the time corresponding to emulated experimental duration, in 1000 simulation steps. The entire ODE simulation can be defined as:

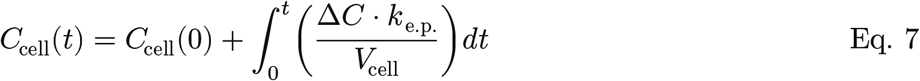

and

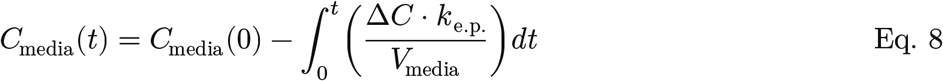

We assumed the initial intracellular H_2_O_2_ concentration to be 50 nM (Imlay, 2013). The concentration gradient between compartments is:

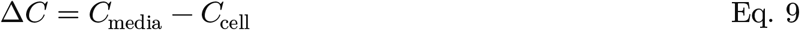

This calculation is specific to each experimental condition and depends on culture density, culture volume, time, and media concentration. The simulation also accounts for the permeability coefficient of H_2_O_2_ (Seaver & Imlay, 2001a):

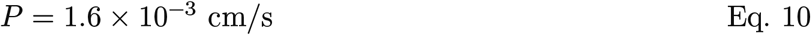

The effective permeability rate (*L* · *h*^−1^) is then defined as:

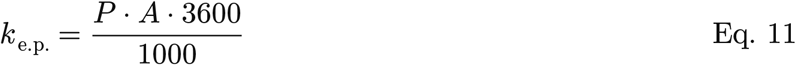

where A is the surface area of the cell (in cm2).

The total H_2_O_2_ uptake (in mmol) was computed as Δ*C*_media_ · *V*_media_ · 1000. Dividing by the total cell number, computed from OD and culture volume, and experimental duration (in h) yielded the average influx (in mmol/ h/cell). Normalization by dry weight (in g) provided the H_2_O_2_ exchange flux in units compatible with our metabolic models.

All constants used in this calculation are included in the Suppl. Information. The average simulated influx was used to set lower and upper bounds on H_2_O_2_ exchange in our *E. coli* metabolic models, deriving input H_2_O_2_ treatment concentrations either from published work (Hashimoto et al., 2005; Jozefczuk et al., 2010; Roth et al., 2022) or estimated from relevant plots (Brandi et al., 1987) using WebPlotDigitizer v4.7 for MacOS.

### Metabolite turnover flux calculation

Some reported analyses and dynamic model constraints rely on tracking metabolite turnover fluxes. Because COBRA models are reaction-centric, where reaction fluxes are the variables being solved for in simulations, metabolites are represented indirectly through the steady-state constraint:

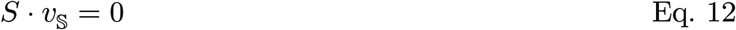

where *S* is the stoichiometric matrix and *v*_S_ is the steady-state reaction flux vector. *S* matrix is indexed as *S*_ij_ where *i* is the index of a specific metabolite (row) and *j* is the index of a specific reaction (column). The metabolite turnover flux values are not directly defined by default but can be mathematically exposed by first transposing the *S* matrix to yield the *S*^*T*^ matrix (reactions as rows, metabolites as columns). This transposed matrix is thus indexed as 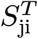 so that *j* and *i* are still reaction and metabolite indices, respectively, and everywhere else in our work. Similarly, the *v*_S_ reaction flux vector, or *v* for short, can be indexed as *v*_*j*_. Thus, 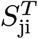 is the stoichiometric coefficient of metabolite i in reaction *j*, and *v*_*j*_ is a steady-state flux of reaction *j*. Illustrated:

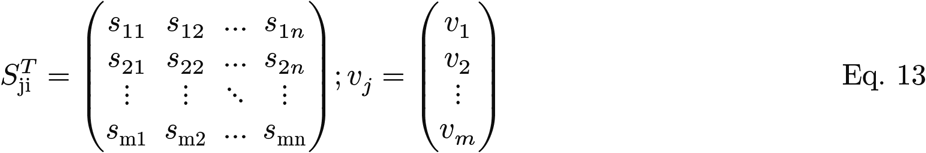

where *m* is the total number of reactions, *n* is the total number of metabolites.

Metabolite turnover flux of metabolite with index *i* can then be expressed mathematically as the sum of products of reaction fluxes *v*_*j*_ and respective metabolite stoichiometric coefficients 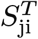 across all reactions that produce the metabolite:

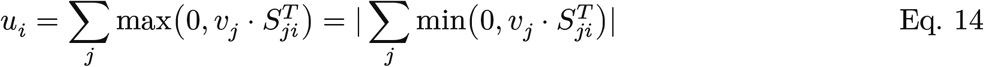

so that *u*_*i*_ is the steady-state metabolite turnover flux of metabolite *i*. Because of the steady-state constraint, the sum of positive metabolite turnover fluxes would always equal the sum of negative (min) metabolite turnover fluxes so only one of them is needed forward.

The metabolite turnover flux calculation, as described here, is normally done for every metabolite as part of the sampling vectors’ analysis. Some of our constraints must leverage the metabolite turnover flux, which means that there is an alternative definition that is done sparsely and dynamically.

### Dynamic constraints

Dynamic constraints allow us to specify relationships between reactions and metabolites that every feasible solution must obey, such as that an SOR must carry flux if the metabolite to be oxidized by ROS in the SOR is produced in the model. Using binary variables to implement conditional constraints as simple logic layers that are not part of the network topology converts the BTC model into a MILP problem. For efficient performance, we implement these constraints using CPLEX’s indicator variables but also show their behavior below through the classic big-*M* variant (Vielma, 2015).

The simplest constraint is the coupling of a binary variable *x*, where *x* ∈ {0, 1}, to a continuous flux variable *y*, where *y* ∈ *R*, such as the flux through a reaction or set of reactions. The binary variable must be equal to 1 if the continuous flux variable is over a threshold *y*_ref_, where *y*_ref_ ∈ *R*. This constraint can be summarized as:

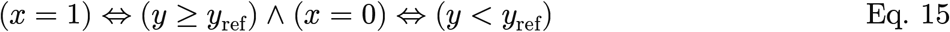

or

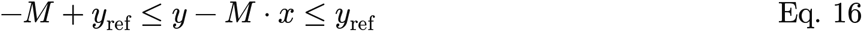

The behavior can also be inverted, when necessary:

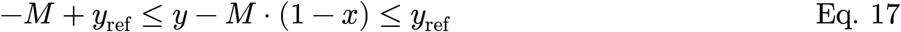

Depending on the application, our big-*M* variables are equal to 1,000, when tracking a continuous flux variable bounded by the same number, or 10,000 when tracking the sum of multiple continuous flux variables.

Since the solver may otherwise violate the upper and lower bounds of this constraint up to the feasibility tolerance, breaking the strictness of our logic, we make this constraint even more restrictive by introducing an additional buffer of feasibility tolerance (*ε*_*f*_):

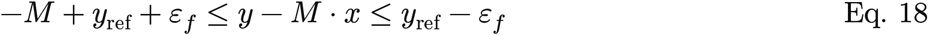

or

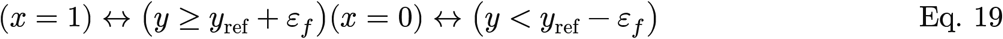

This modification also effectively makes the non-zero flux threshold 2 · *ε*_*f*_ throughout our models. This binarization function can be summarized as:

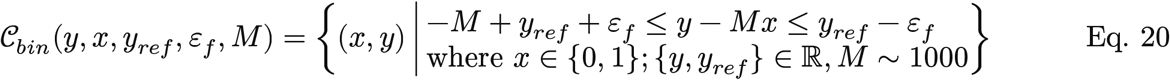

or

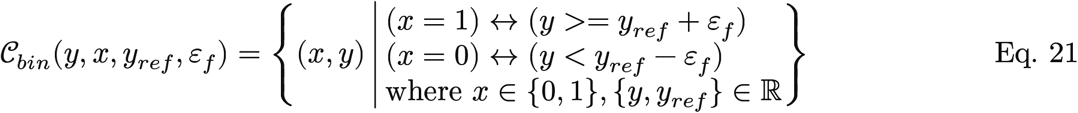

Two or more continuous flux variables may share the same binary variable, coupling their behavior always. An alternative is to have a separate binary variable coupled to continuous variables:

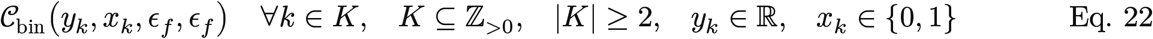

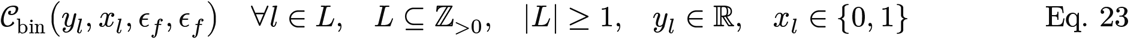

where *K* is a set of inputs and *L* is a set of outputs, which then allows conditional coupling through the binary variables via the use of an AND logic gate. This logic circuit ensures that all the binary outputs have a value of 1 when all binary inputs also have a value of 1 (and same in the opposite direction):

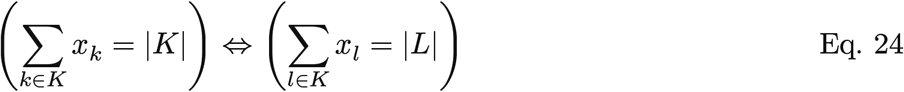

which is equivalent to:

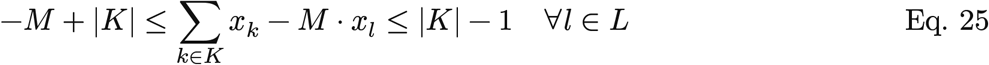

This logic circuit allows us to binarize an SOR flux, in addition to binarizing the metabolite turnover fluxes of its two reactants. The AND gate is then used to enforce the usage of the SOR when both its reactants are present.

Because it is necessary for the AND gates, metabolite turnover fluxes (defined previously) can be dynamically exposed in a sparse fashion. Given the stoichiometric *S* matrix and steady-state reaction flux vector *v*_S_ or *v*, the canonical steady-state constraint is defined as:

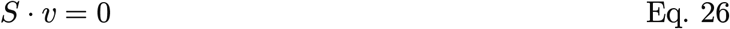

Metabolite turnover flux can be dynamically expressed with the help of the transposed *S* matrix 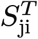 so that the row index corresponds to reactions *j* and column index corresponds to metabolites *i* for consistency. Broadcasting of the product of the *S*^*T*^ matrix columns and the reaction flux vector *v* as a matrix *H* represents the production and consumption fluxes for each metabolite by every reaction:

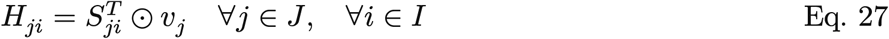

where *J* (rows) is the set of all reaction indices; *I* (columns) is the set of metabolite indices participating in SORs, which makes the *H* matrix a sparse version of *S*^*T*^ because the operations below are expensive to do dynamically. A mathematical circuit was then be created to dynamically constrain the *H* matrix, yielding the *P* matrix, in which negative values are set to 0 using a binary variable that tracks the net contribution of each reaction to each metabolite of interest:

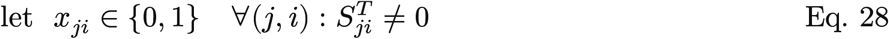

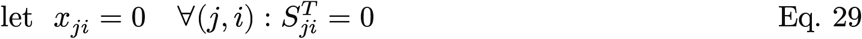

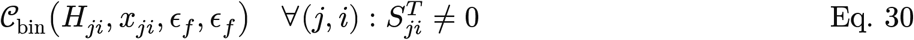

The absolute value of *H*_ji_ can then be dynamically computed as an MILP problem. To do this we define a new matrix to hold the absolute values:

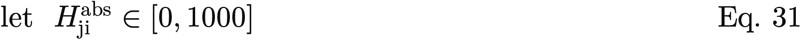

and couple the two *H* matrices:

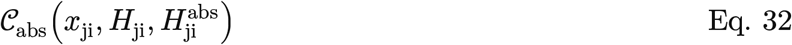

which enforces the conditional behavior:

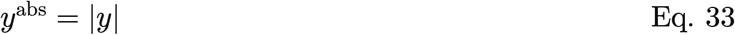

Implementing this circuit consists of four equations. The first two constrain the lower limit on *y*^abs^ to be |*y*| when *y* is positive and negative, respectively:

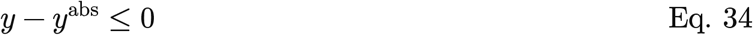

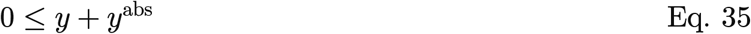

The second set of equations constrain the upper limit on *y*^abs^ to be y when y is positive and negative, respectively:

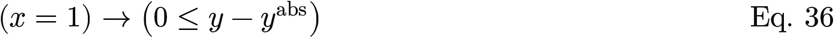

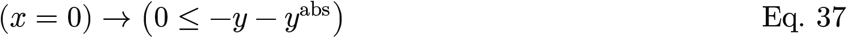

This second set of equations can also be expressed via big-*M*, which needs to be larger than 2000 in this case:

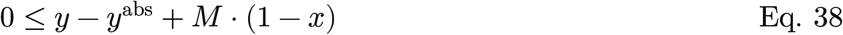

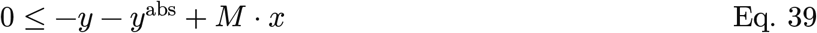

Yielding the constraint:

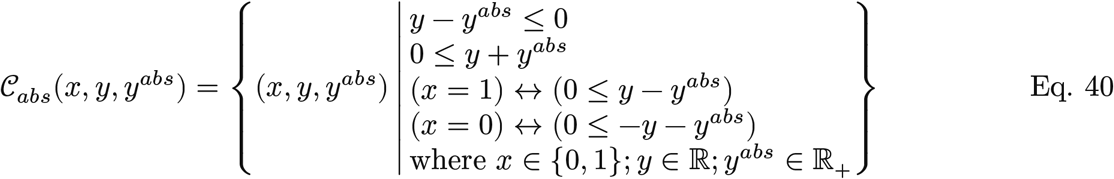

or

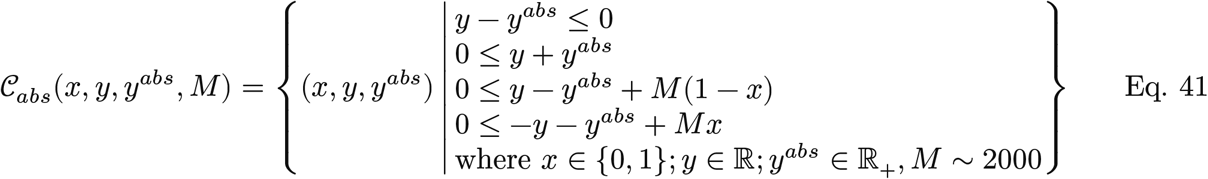

The 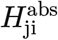 constrained matrix allows the solver to easily compute *P*_ji_ as:

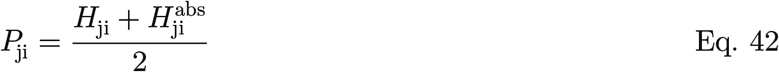

*P*_ji_ can then be used to define a metabolite turnover flux vector, containing the metabolites of interest, by aggregating the net positive contributions from reactions:

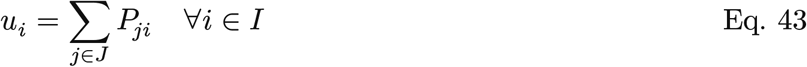

The sparse metabolite turnover flux vector can then be referenced by AND gates to enforce usage of an SOR when all its reactants have non-zero turnover. These constraints enable control of SOR usage and rates. Controlling reaction rates consists of a separate constraint system detailed in the Suppl. Methods.

### Optimization and Flux sampling

All our optimizations were performed with the CPLEX solver. Details on its configuration can be found in the Suppl. Methods section.

For sampling we used a modified version of GapSplit (Keaty & Jensen, 2020) supporting multi-core parallelization and sample quality control to maximize diversity across the solution space. Warm-start vectors and target ranges for core reactions were generated using parallelizable COBRApy FVA without biomass constraints; because BTC is a MILP model, these were instead calculated from the corresponding base model under each culture condition. Random target-gap selection generated stochastic Markov chains, with independent chains serving as biological replicates. At least 2,500 random-gap samples per replicate were required for convergence, so 5,000–10,000 samples were generated before quality-control filtering. Model-specific sampling parameters, HPC implementation, and filtering criteria are detailed in the Suppl. Methods.

### Survivor, persister, and grower classes

Stochastic flux samples can be classified as survivors and non-survivors. Non-survivors are feasible solutions with ATPM (ATP maintenance) flux less than 3.15, derived from the ATPM lower bound in iJO1366. Sampled ATPM fluxes naturally form a bimodal distribution, well-separated by a 3.15 threshold. Survivors are divided into persisters and growers. Growers have a biomass flux greater than twice the feasibility tolerance, the *de facto* non-zero flux threshold for our models. Samples simulated in LB medium in the BTC and BT models showed a distinct transition in biomass flux around 80% of maximum biomass (**Figure S5**), leading us to classify the top 20% of growers based on biomass value in some analyses to distinguish near-optimum growth metabolic mechanisms from those of the remainder (bottom 80%) of most growers.

### Selecting for metabolic hysteresis

To identify H_2_O_2_-exposed solutions consistent with metabolic hysteresis, we defined a pre-treatment metabolic range based on the flux vectors of untreated growers in the top 20% of biomass production. These vectors were averaged to obtain a single untreated centroid representing the common pre-culture state. We calculated Euclidean distances between each sampled treatment vector and this centroid, retaining only solutions whose distances fell within the minimum and maximum distances observed among the untreated growers. This range was used to identify treatment solutions that could plausibly arise through hysteresis from the common pre-treatment state. This approach was described in more detail in the Suppl. Methods section.

### Metabolomics data processing

Raw metabolomics peak height data from a previous study (Jozefczuk et al., 2010) was scaled to sample optical density. For each metabolite in at each time point, the average was taken first across technical replicates, then across biological replicates. Biological replicates without three technical replicates were discarded from the averaging. Data from 0 min and 10 min post-treatment oxidative stress samples, corresponding to time points 3 and 4 in the published experiment (Jozefczuk et al., 2010), were selected as the control and treated conditions, respectively. The treatment consisted of extracellular exposure to 200 μg/mL H_2_O_2_, which corresponds to 5.88 mM H_2_O_2_. Namespace matching was then performed between the names of metabolites as given in supplemental information in the prior study (Jozefczuk et al., 2010) and metabolite names in our BTC model to obtain the intersection of metabolites in both the model and the metabolomics dataset (Suppl. Information file). The validation dataset metabolites were aggregated by summing synonymous entries identified as follows based on careful review of names and descriptors as previously published. First, unambiguous matches were identified, unidentified compounds from the metabolomics (unknowns) were excluded, and major and minor metabolite peaks were treated as the same metabolite. Incompletely specified metabolomics results including “similar to” in their descriptions were then mapped to a BTC model metabolite only if that metabolite was not already unambiguously matched. Linear and cyclic isomers were treated as the same metabolite when only one form is represented in the BTC model. Entries from the metabolomics describing two or more metabolites are validated jointly if both are in the BTC model. Modified metabolite forms (e.g. trimethyl lysine) are mapped to their respective model entries if one exists, otherwise counted toward the unmodified form (e.g. lysine). Similarly, the predicted metabolite flux data had to be filtered before comparing to avoid thermodynamically infeasible loops (see Supplementary Methods).

### Metabolomics evaluation

The intersection of the processed and filtered metabolites measured in the previously published metabolomics study (Jozefczuk et al., 2010) and simulated and loop-corrected metabolite turnover fluxes was used to evaluate how relatively well individual sampled flux vectors and median fluxes aggregated across samples from our models recapitulated *in vitro* oxidative stress metabolic phenotype. Because the metabolomics and simulated flux datasets represent abundance and turnover, respectively, they are not directly comparable. Instead, we compare pairwise metabolite ratios between these datasets to capture the relative distribution of metabolic activity across metabolite nodes in the network. These ratios are represented in a two-dimensional symmetric matrix with rows and columns representing the same list of metabolites. The upper triangular matrix from the simulated and measured datasets are then extracted, flattened into a one-dimensional vector, and their cosine similarity is calculated. This analysis was performed for both H_2_O_2_-treated and untreated conditions and across simulated survivor, grower, and persister classes.

## Acknowledgements

This research was supported in part by the National Institute of General Medical Sciences of the National Institutes of Health under award number R35GM151236 to RLC.

## Supplementary Methods

### Solver configuration

All optimizations were performed using IBM’s CPLEX solver (v22.1.0.0) through COBRApy, available from PyPi as *cobra* (v0.26.3), using *optlang* (v1.8.3) to interface with the solver. The optimality, integrality and feasibility tolerances were set to 1 × 10^−9^. We enabled the solver maximum multithreading on Einstein’s high performance computing cluster (HPC) with 32 CPUs per node. Our computer cluster did not support hyperthreading; each thread was handled by a separate CPU. We allowed 8 GB of memory per CPU up to the total node memory per optimization. We also used deterministic parallelism for all CPLEX optimizations, unless stated otherwise, to ensure that re-optimization of the same problem yields identical results. We disabled pre-solving, not beneficial to our MILP optimizations. Our model constraints require stricter adherence to variable bounds, which we enabled with the *numerical emphasis switch* (see IBM documentation). We gave higher branching priority to core reactions of the model through a priority input file of a list of tuples representing each variable’s name, its importance, and preferred direction. The variables in this list include the base model reactions from iJO1366, with an importance of 2, in addition to the scaling variables from the ROS-chemistry expansion with an importance of 1. All preferred directions were set to 0. Each optimization was given 2000 s to accommodate that most biomass optimizations needed between 20 and 30 min in our BTC model. Rare cases that the solver returned infeasible or no solution were resolved by re-running the optimization a up to four times, usually less than 2 h total time for optimization. Stochastic flux sample generation was not given extra attempts, and each optimization had a 300 s timeout. To conserve system storage, infeasible or timed-out samples were not saved.

### Flux sampling

GapSplit target gaps were selected randomly, with each new target continuing from the previously closed gap, thereby representing stochastic transitions through the solution space. Independent realizations of this process were treated as biological replicates. Sampling parameters differed among model formulations: models B and BT used a primary tolerance of 10^−3^ around the target-gap center and a secondary-gap fraction of 5%, whereas BTC required a primary tolerance of 0.1 and a secondary fraction of 10^−5^, effectively eliminating secondary targets.

Because Einstein HPC restrictions prevented parallelization within individual Markov chains, samples were generated consecutively while relying on CPLEX multithreading for the underlying optimization. Each Markov-chain job was limited to 24 h, which was sufficient to obtain the required number of samples. Preliminary benchmarking indicated that convergence required at least 2,500 random-gap samples per replicate (data not shown); therefore, 5,000–10,000 samples were collected before filtering.

Quality control excluded samples violating steady-state requirements, ROS-reaction AND-gate logic, binary constraints, death accounting, or reaction bounds. Biomass constraints were intentionally omitted during sampling, with differential biomass instead incorporated by classifying flux samples during downstream analysis. H_2_O_2_ exchange upper and lower bounds were set to the same value, dependent on experiment and condition and calculated through our ODEs, in the interest of improving compute time.

### Selecting for metabolic hysteresis

The pre-treatment metabolic range was constructed from untreated growers with biomass flux in the upper 20% of the untreated population, based on the assumption that permissive conditions favor optimization of biomass production (Goelzer et al., 2015; Harcombe et al., 2014; Molenaar et al., 2009; Schuetz et al., 2012). The flux vectors of these growers were averaged component-wise to obtain their centroid.

Because the emulated treated conditions do not represent descendants of the untreated culture, but instead originate independently from a shared pre-culture state, we did not use individual untreated samples as references. The centroid therefore represents the common starting metabolic state, conceptually analogous to a single founding colony.

For each untreated top 20% grower, its Euclidean distance from the centroid was calculated. The minimum and maximum of these distances defined the lower and upper boundaries of the pre-treatment metabolic range. Euclidean distance was then calculated between every sampled treatment vector and the untreated centroid. Treatment vectors falling outside these boundaries were discarded because their metabolic displacement from the common starting state exceeded the range observed among untreated growers and was therefore considered inconsistent with the modeled hysteretic transition.

### Accounting for flux from thermodynamically infeasible loops

After selecting flux sampling data that represent hysteresis as described above, samples were analyzed to identify reaction fluxes driven by thermodynamically infeasible loops. Because the previously published CycleFreeFlux method (Desouki et al., 2015) cannot be used with our models due to their dynamic constraints, we developed a custom approach instead.

A hallmark of thermodynamically infeasible loops is reaction fluxes equal to or approximating the default numerical bounds (± 1000); any reaction that can carry flux with |*v*| ≥ 999 is expected to participate in such a loop. To identify reaction participating in loops from the base model, we closed all exchange reactions, removed all dynamic constraints, and set the ATP maintenance reaction flux lower bound to zero before performing flux variability analysis (FVA) (Mahadevan & Schilling, 2003) to determine the minimum and maximum fluxes possible for each reaction. Any reactions carrying near-maximum flux under these minimal conditions were flagged as participating in loops, and these reactions and their metabolites were not used in downstream analysis.

As another correction for possible looping flux contribution to metabolite turnover calculations, we determined the maximum possible turnover flux for each metabolite. In a separate model, we created a demand reaction for each metabolite and maximized flux through this demand reaction set as the objective in an FBA simulation. These results indicate the maximum possible metabolite turnover flux that does not incorporate flux through thermodynamically infeasible loops. We used these values at metabolite-specific ceilings for identifying additional reactions and metabolites to exclude.

### ROS consumption as a function of rate constants

When SORs are active under the AND gate dynamic constraints, their fluxes must be a function of their reaction rate constants and the degree of ROS exposure, represented as a scaling variable for each ROS. Each SOR reactant is similarly represented by an independent scaling variable, and every SOR flux is independently determined by each of its two reactants’ scaling variables, which are thus said to be *coupled* by default to support the flux as long as both reactants are sufficiently available. If a reactant’s availability is limiting for SOR fluxes, the respective scaling variables prevent metabolite and ROS overdemand going beyond mathematical limits by decoupling those metabolite scaling variables from the set of SOR fluxes involving these metabolites, and residual ROS availability is thus distributed across the remainder of SOR reactants. Decoupling either one of the reactants therefore results in the respective SOR reaction fluxes being informed by only the other reactant’s scaling variable. In rare circumstances double-decoupling may occur for both reactants, causing the SOR to usually take on a value of two times the solver feasibility tolerance. To prevent solver cheating by arbitrarily increasing scaling variables and forcing double decoupling and complete freedom in most circumstances, a constraint was imposed requiring that ≥ 98% of each decoupled reactant be consumed by SORs. Because requiring that a high percentage of oxidizable metabolites are consumed by SORs competes with biological objectives like biomass, the solver only decouples metabolite scaling variables when no other feasible solution exists. Implementing this mathematical circuit requires explicit treatment of each SOR reactant and each rate-constrained SOR.

To encode the handling of metabolites participating in SORs, we hardcode a negative stoichiometric matrix *N* that includes only the originally negative coefficients from *S*^*T*^ matrix which is a transposed *S* matrix where rows (*j*) are now indexed by reactions and columns (*i*) are now indexed by metabolites:

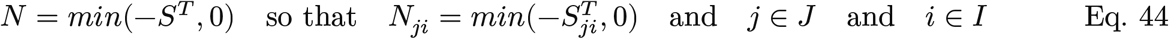

where *J* is a set of all SOR reactions, and *I* is a set of all their reactants. We then multiply the matrix *N* element-wise by the reaction flux vector *v*, thus broadcasting *v* across the corresponding dimension of *N*:

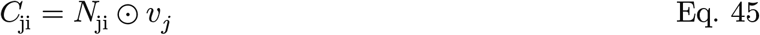

which is the metabolite consumption matrix *C* expressed as positive values. Then the SOR reactant availability flux vector *a*_*i*_ is computed from the metabolite flux vector *u*_*i*_ as:

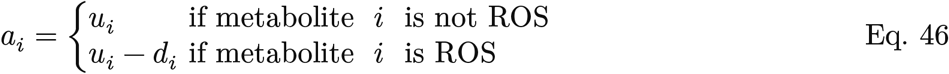

In the case of ROS, this flux available to SORs is corrected with flux consumed by the macromolecule demand reactions whose flux is the same as their only reactant’s, summarized in flux vector *d*_*i*_. We then define a binary overdemand variable for each SOR reactant:

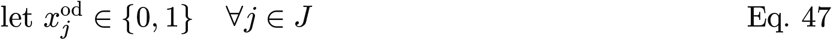

Another variable is then defined for the requested reaction fluxes, those determined by the reaction rate constant *k* and the scaling variable *s*. This matrix *K* has the same shape as *N*:

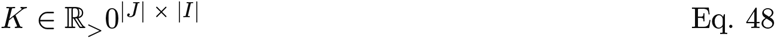

Coupling of the binary overdemand variables and matrix *K* is then invoked by tracking the difference between what flux is available and what has been requested by all SORs for a given reactant:

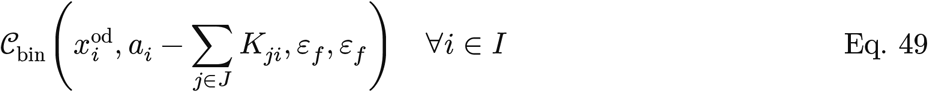

Finally, the 98% metabolite overdemand rule is enforced:

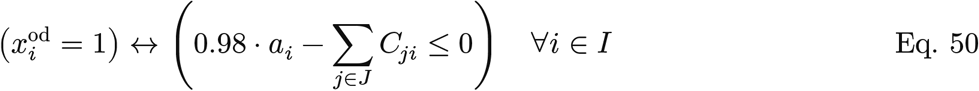

To enforce SOR flux scaling, we hardcode a vector *k*_*j*_ containing all of the SOR rate constants. We also define a vector *s*_*i*_ that holds the solver-determined scaling variables for each SOR reactant *i*. For each SOR with index *j* we define three sets of equations, controlled by the binary reaction usage variable *x*_*j*_ of that reaction (see main Methods).

The first set of equations determines the SOR-requested flux from the oxidant ROS as *K*_io_, whose scaling variable is *s*_*o*_ ∈ ℝ_+_:

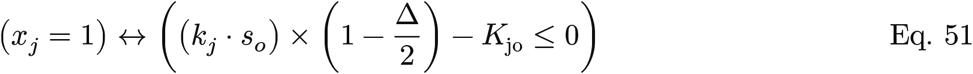

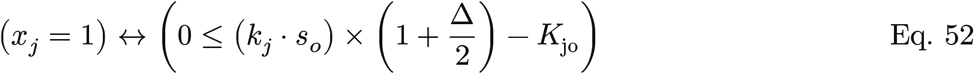

A second set of equations determines the SOR-requested flux from the reactant being oxidized as *K*_ir_, whose scaling variable is *s*_*r*_ ∈ ℝ_+_:

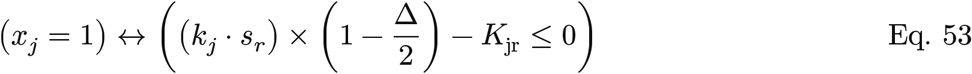

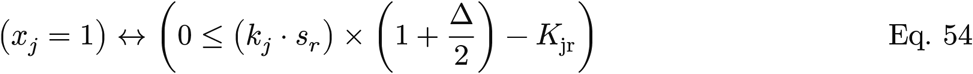

Note that Δ is a constant providing buffer around the requested reaction flux, set at 0.05 by default. When the given SOR is not active, a third set of equations takes precedence, ensuring that the given SOR requests no metabolite flux:

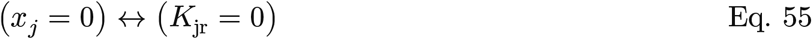

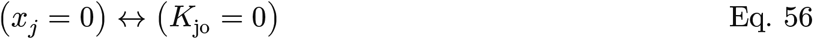

With these constraints in place, the solver will find a set of values for *s*_*i*_ that satisfy all constraints. Finally, the requested metabolite flux must be connected to the SOR flux. This also handles decoupling of SORs from a given metabolite scaling. We use the vector of metabolite overdemand binary variables 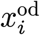 defined earlier so that for each SOR *i*:

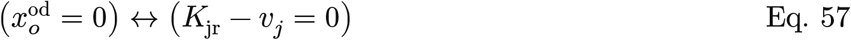

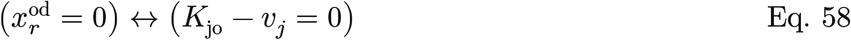

Notice that 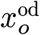 controls the coupling of *K*_jr_ with *v*_*j*_, and 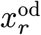 informs the coupling of *K*_jo_ with *v* _*j*_. So, if the oxidant ROS is in overdemand the oxidizable reactant will not determine the SOR reaction flux, and if the oxidizable reactant is in overdemand then the ROS will not determine the SOR reaction flux.

## Supplementary Figures

**FIGURE S1.**
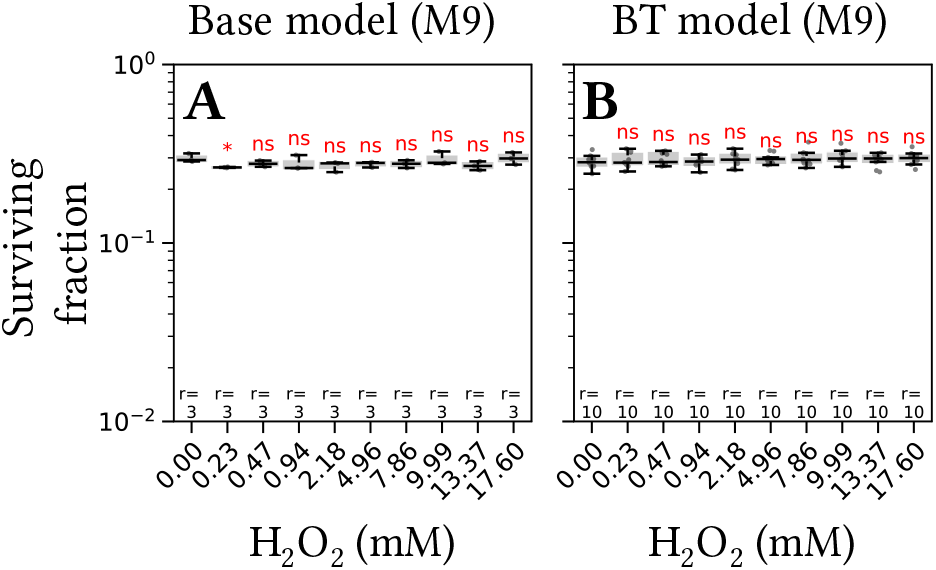
Expanded ROS topology alone does not reproduce bimodal killing by H_2_O_2_ in genomescale models. M9 versions of the Base **(A)** and BT **(B)** models were stochastically sampled with GapSplit, emulating the same experimental setup as Brandi et al. (1987). Flux vectors were filtered for enforcing hysteresis, and survivor fractions are shown. Significance by t-test (not corrected given small n): ns, p≥0.05; *, p<0.05.

**FIGURE S2.**
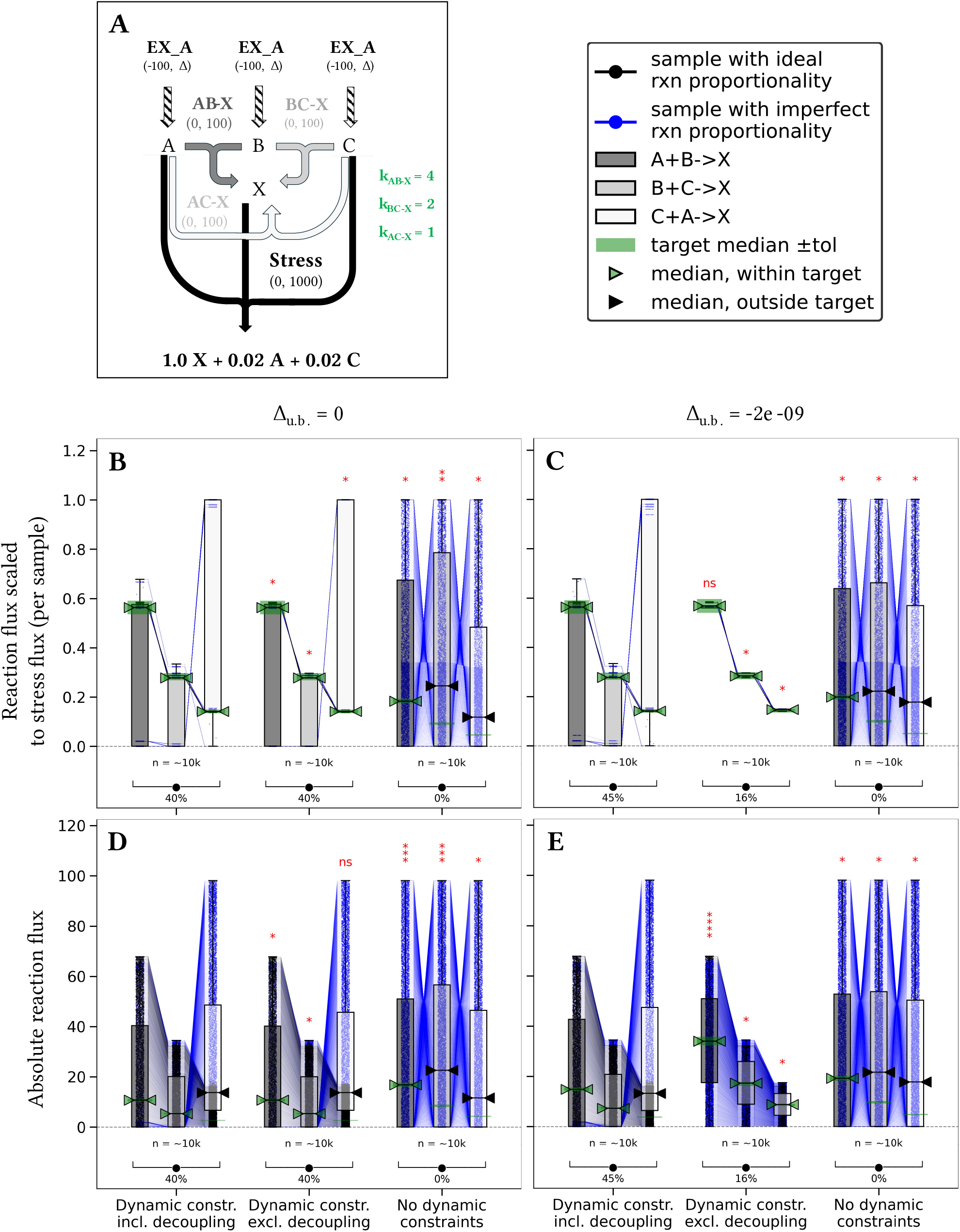
Numeric validation of the Compulsion, Proportionality, and Decoupling constraint system. **(A)** MILP representation of the SOR subset from Figure 2. Metabolites A, B, C stand in for natural products (some possibly ROS); X is their interaction byproduct via A+B→X, B+C→X, or C+A→X (grey, stoichiometry −1 per reactant). The Stress demand reaction (black) emulates distinct shadow prices, which, with rate constants (green), set pull through the exchanges (striped arrows). Exchanges are flexible (**Δ**=0) or forced (**Δ**= 2 × 10^−9^ see Methods); forced exchanges rely on AND-gate constraints to engage all three SORs – the normal use case. **(B-E)** Three models per pabel: dynamic constraints with decoupling, without decoupling, and none (dynamic constraints = AND gates + proportionality). Fluxes shown as modified box plots (legend upper right); small dots are individual samples, lines connect the same sample across its three reactions. Black dots/lines mark ideal samples (all three fluxes perfectly proportional); blue mark samples with one or more reactions out of lockstep (due to exchange rate and/or decoupling). Green boxes give expected medians relative to A+B→X; green/black triangles indicate medians inside/outside those boxes. Sample counts and ideal-sample fractions appear below each set. **(B, C)** Flexible and forced exchanges, respectively, scaled to Stress per solution. **(D, E)** Raw fluxes over the full sampling range. Comparisons are per reaction (Mann-Whitney, Bonferroni-corrected, dynamic+decoupling as control): ns, p≥0.05; ** p<0.01; ***, p<0.001; ****, p<0.0001.

**FIGURE S3.**
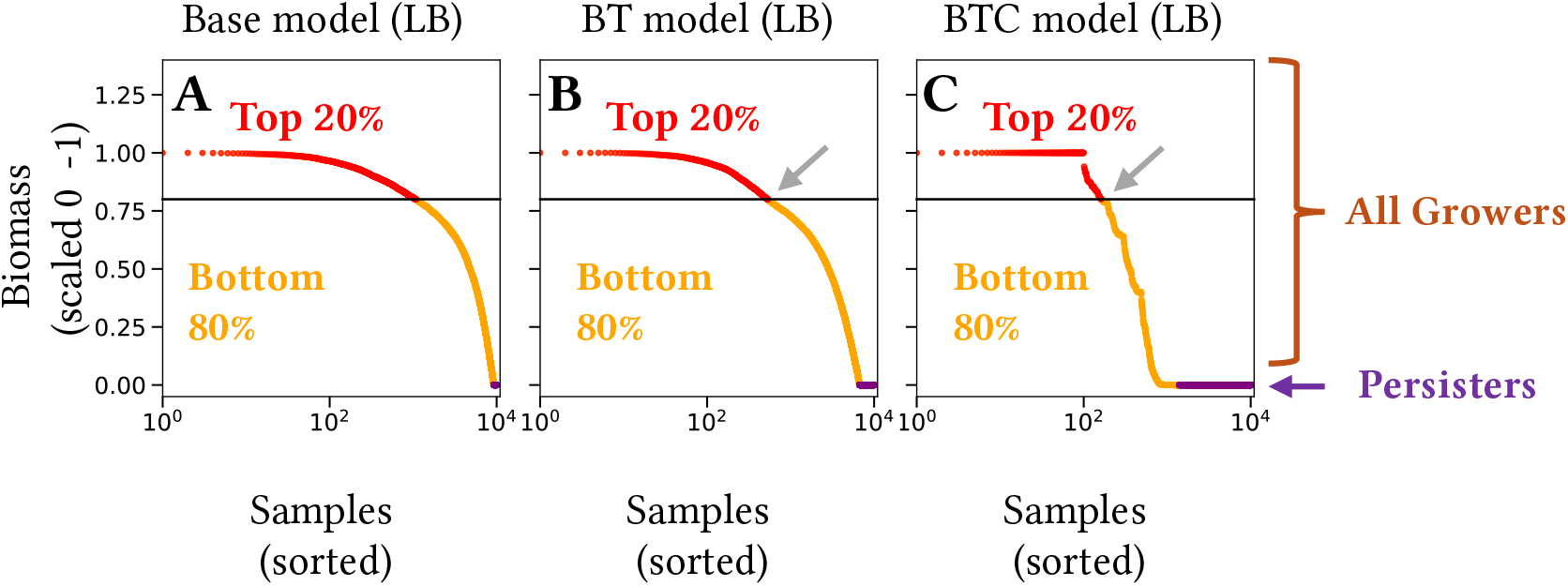
Identification of distinct survivor subsets in LB medium. The Base, BT and BTC genome-scale MILP models in LB were stochastically sampled with GapSplit, emulating the experiment of Hashimoto et al. (2005) (no H_2_O_2_ treatment). Replicates were pooled per model; panels **A-C** show the fraction of maximum biomass for Base, BT, and BTC, Black lines mark 80% of each sample group’s maximum biomass – the threshold nearest the distinct mode shift in BT/BTC (grey arrow). This subdivides growers (net positive biomass) into Top 20% (red) and Bottom 80% (orange) of maximum biomass; persisters (survivors with zero biomass) are shown in purple.

**FIGURE S4.**
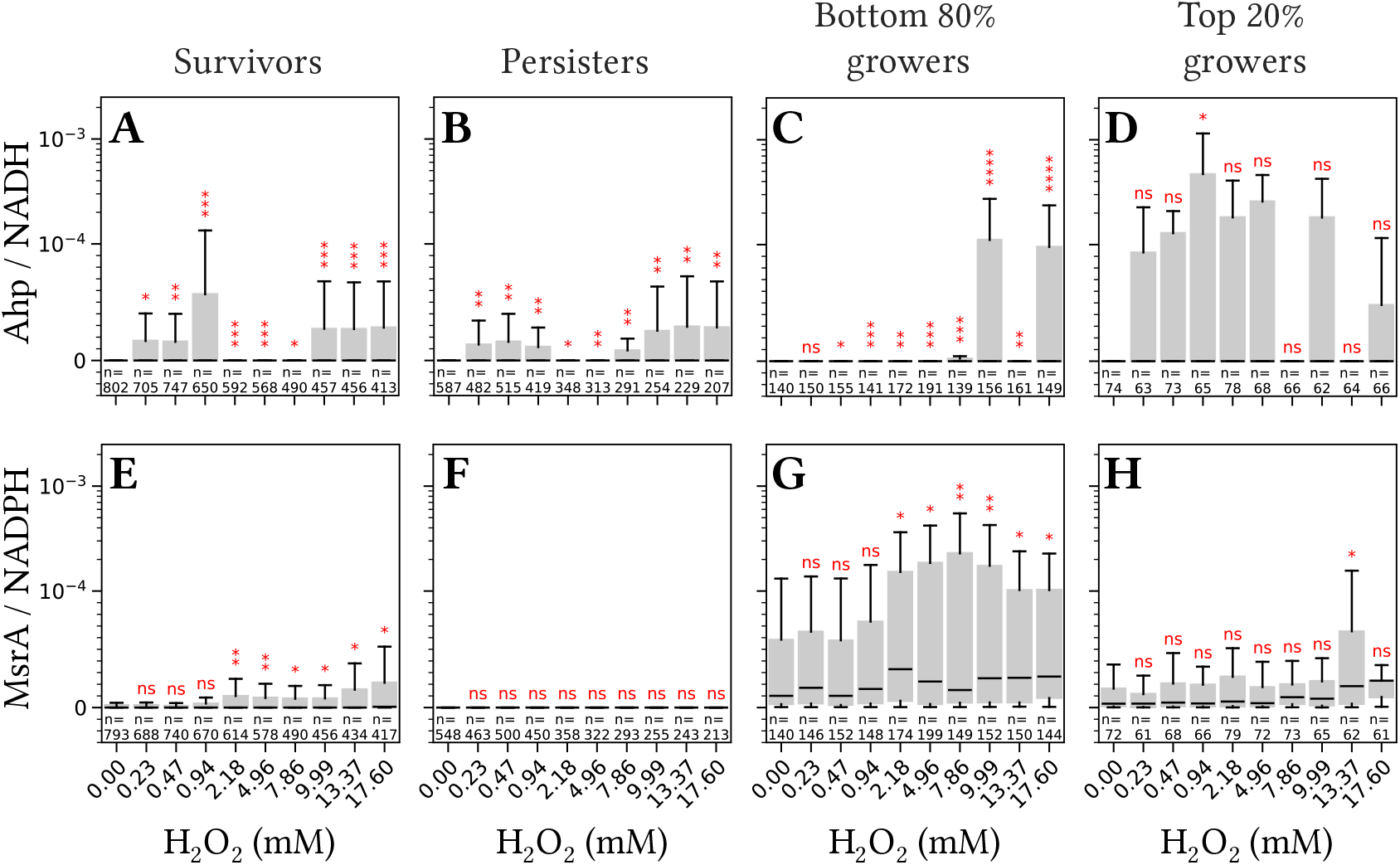
Persisters rely almost exclusively on Ahp, whose activity rises with recovery onset. BTC (M9) samples were filtered by the hysteresis vector distance-based cutoff criterion (see Methods) and grouped into survivors, persisters, bottom 80%, top 20% of growers (by biomass). Ahp and MsrA fluxes were scaled by the median NADH and NADPH turnover fluxes per sample. **(A-D)** Scaled Ahp use in survivos, persisters, bottom 80%, and top 20% of growers. **(E-H)** MsrA use, same order. Significance by t-test (uncorrected): ns p≥0.05, * p<0.01, *** p<0.001, **** p<0.0001.

**FIGURE S5.**
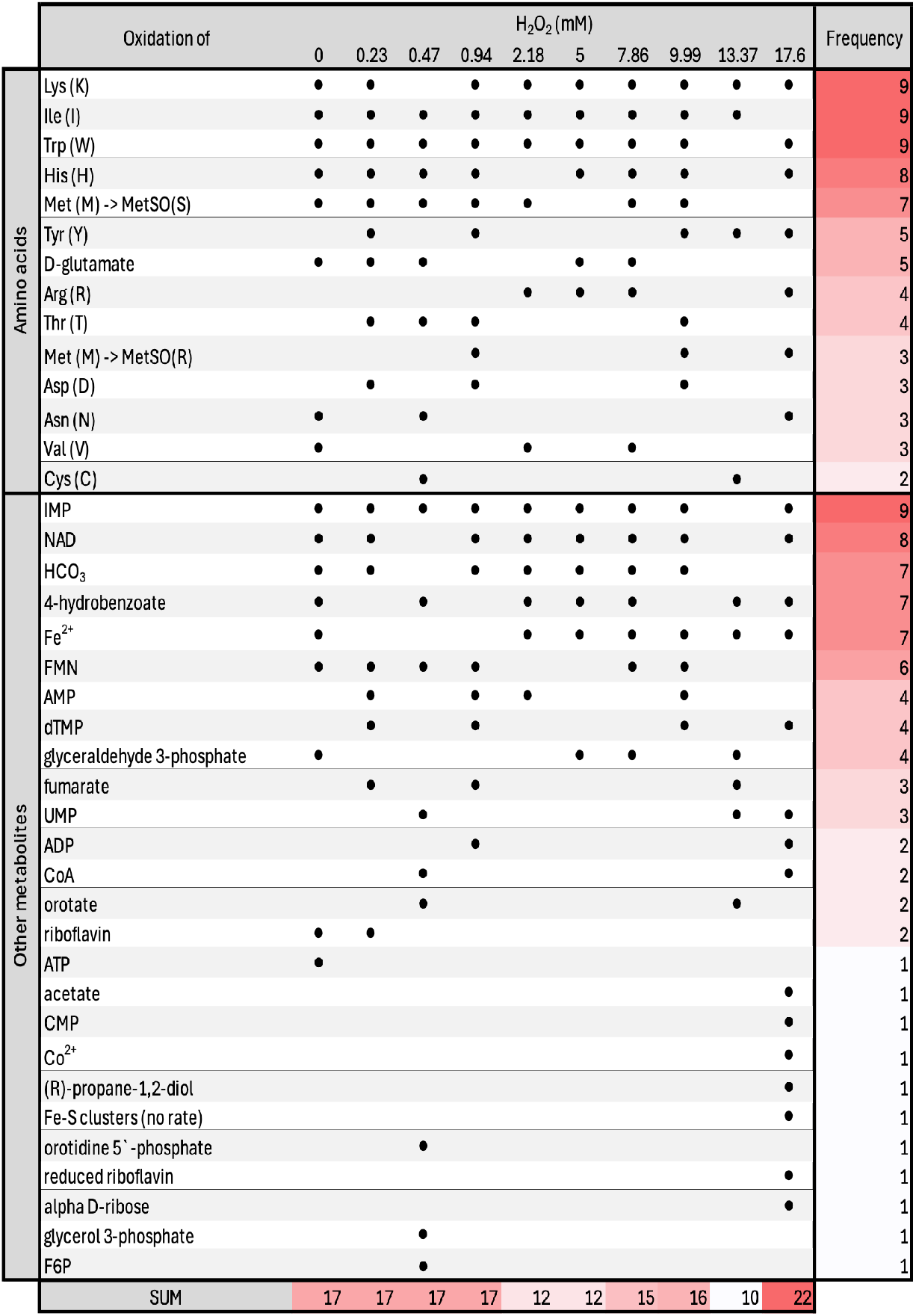
Frequency of SORs by H_2_O_2_ concentration. The SOR PCA from Figure 5B was repeated per H_2_O_2_ concentration; 95%-filtered variable loadings associated with growers were counted and shown as oxidation modalities. • indicates oxidation of a given metabolite at that [H_2_O_2_].

**FIGURE S6.**
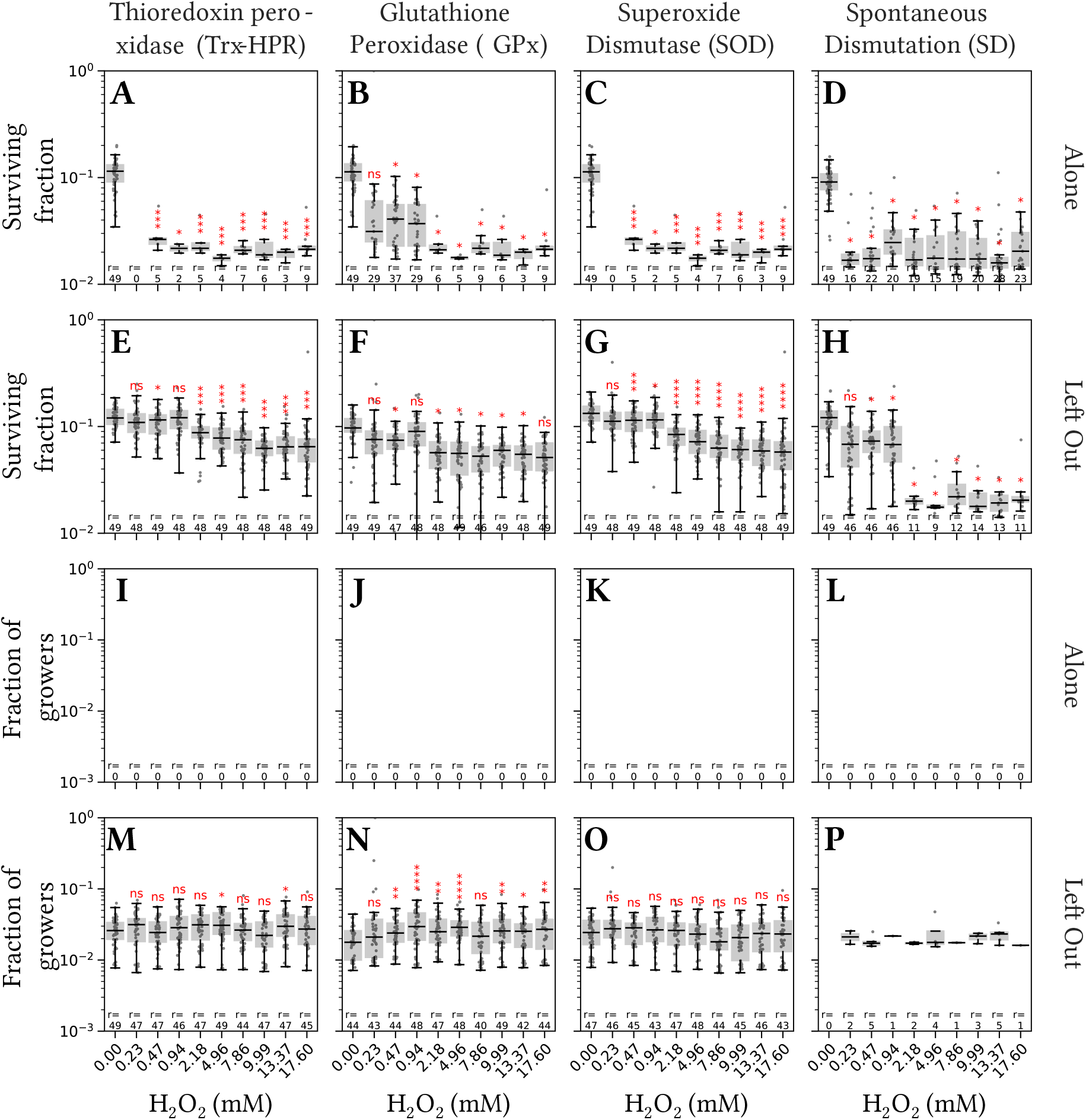
Baseline survival requires both glutathione peroxidase and spontaneous dismutation. The Trx-HPR + GPx + SOD + SD combination was dissected as before to identify which reactions underlie the observed baseline survival in Figure 6. **(A-D)** Survival when each is solely available. **(E-H)** Survival when each is the only one disabled. **(I-L)** Grower fraction when each is solely used. **(M-P)** Grower fraction when each is exclusively used. Significance by t-test (uncorrected): ns p≥0.05, * p<0.05, ** p<0.01, *** p<0.001, **** p<0.0001.

**FIGURE S7.**
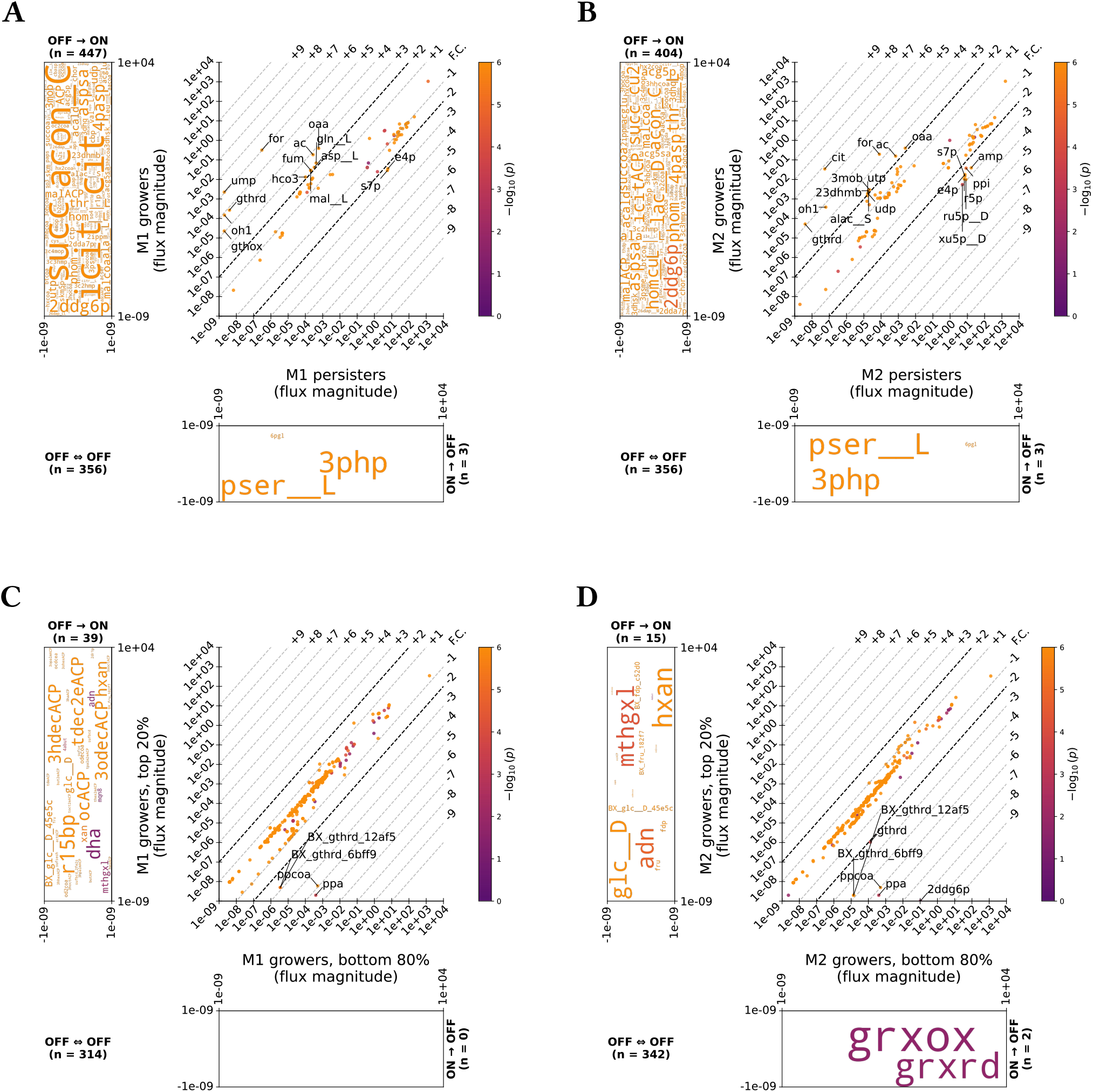
Persisters generate NADH through 3-Phospho-D-glycerate while top growers route PPP flux through EntnerDoudoroff pathway and back into glycolysis. Flux vectors from BTC (M9) were filtered by the hysteresis vector distance-based cutoff criterion (see Methods); metabolite turnover was computed for persisters, all growers, and top 20%/bottom 80% of growers by biomass (see Figure S3). Samples were split by H_2_O_2_ range before comparison. Samples were split by H_2_O_2_ range before comparison: Mode 1 (M1, x-axis) is 0, 0.23, 0.47, 0.94, or [0, 2 mM); Mode 2 (M2, y-axis) is 2.18, 4.96, 7.86, 9.99, or [2, 10 mM). Flux magnitudes are expressed in mmol/gDW/h. **(A, B)** All growers versus persisters, Mode 1 and Mode 2. **(C, D)** Bottom 80% versus top 20% of growers, Mode 1 and Mode 2. Plot areas are colored by p-value (color bar); only metabolites passing the Mann-Whitney significance threshold (p ≤ 0.05) are shown. Cytoplasmic metabolites were used, except those lacking a cytoplasmic version. The scatter shows per-metabolite magnitude change between modes; the center diagonal marks no change, thick dashed lines mark changes within two orders of magnitude (metabolites beyond are labeled). Wordclouds show metabolites transitioning ON →OFF (bottom) and OFF→ON (left) from M1 to M2, where ON/OFF is defined by a single flux value higher than the feasibility tolerance (1 × 10^−9^); word size reflects relative ON-state flux. The lower-left corner shows metabolites OFF in both modes.

## References

Addorisio, S. R., Shteynberg, R., Dasilva, M., Mixon, J., Mucciarone, K., Vu, L., Arsenault, K., Briand, V., Parker, S., Smith, S., Vise, C., Pina, C., & Laranjo, L. (2022). Oxidative Stress Response in Bacteria: A Review. Fine Focus, 8(1), 36–46. 10.33043/FF.8.1.36-46

Amato, S. M., Fazen, C. H., Henry, T. C., Mok, W. W. K., Orman, M. A., Sandvik, E. L., Volzing, K. G., & Brynildsen,M. P. (2014). The Role of Metabolism in Bacterial Persistence. Frontiers in Microbiology, 5. 10.3389/fmicb.2014.00070

Ambrus, A., Torocsik, B., Tretter, L., Ozohanics, O., & Adam-Vizi, V. (2011). Stimulation of Reactive Oxygen Species Generation by Disease-Causing Mutations of Lipoamide Dehydrogenase. Human Molecular Genetics, 20(15), 2984–2995. 10.1093/hmg/ddr202

Amici, A., Levine, R. L., Tsai, L., & Stadtman, E. R. (1989). Conversion of Amino Acid Residues in Proteins and Amino Acid Homopolymers to Carbonyl Derivatives by Metal-Catalyzed Oxidation Reactions. Journal of Biological Chemistry, 264(6), 3341–3346. 10.1016/S0021-9258(18)94071-8

Asad, N. R., Asad, L. M. B. O., Almeida, C. E. B. D., Felzenszwalb, I., Cabral-Neto, J. B., & Leitão, A. C. (2004). Several Pathways of Hydrogen Peroxide Action That Damage the E. Coli Genome. Genetics and Molecular Biology, 27(2), 291–303. 10.1590/S1415-47572004000200026

Asada, K. (1999). THE WATER-WATER CYCLE IN CHLOROPLASTS: Scavenging of Active Oxygens and Dissipation of Excess Photons. Annual Review of Plant Physiology and Plant Molecular Biology, 50(1), 601–639. 10.1146/annurev.arplant.50.1.601

Atack, J. M., & Kelly, D. J. (2008). Contribution of the Stereospecific Methionine Sulphoxide Reductases MsrA and MsrB to Oxidative and Nitrosative Stress Resistance in the Food-Borne Pathogen Campylobacter Jejuni. Microbiology, 154(8), 2219–2230. 10.1099/mic.0.2008/019711-0

Baffy, G. (2010). Uncoupling Protein-2 and Cancer. Mitochondrion, 10(3), 243–252. 10.1016/j.mito.2009.12.143

Balaban, N. Q., Merrin, J., Chait, R., Kowalik, L., & Leibler, S. (2004). Bacterial Persistence as a Phenotypic Switch.Science, 305(5690), 1622–1625. 10.1126/science.1099390

Bao, X., Yang, J., De Soet, J., Liu, H., Gao, X., Van Loveren, C., & Deng, D. (2017). Factors Influencing the Competition between Streptococcus Oligofermentans and Streptococcus Mutans in Dual-Species Biofilms. Caries Research, 51(5), 507–514. 10.1159/000479044

Bertrand, R. L. (2019). Lag Phase Is a Dynamic, Organized, Adaptive, and Evolvable Period That Prepares Bacteria for Cell Division. Journal of Bacteriology, 201(7). 10.1128/JB.00697-18

Bhalla, P., Rengaswamy, R., Karunagaran, D., Suraishkumar, G. K., & Sahoo, S. (2022). Metabolic Modeling of Host–Microbe Interactions for Therapeutics in Colorectal Cancer. Npj Systems Biology and Applications, 8(1), 1. 10.1038/s41540-021-00210-9

Bielski, B. H. J., Cabelli, D. E., Arudi, R. L., & Ross, A. B. (1985). Reactivity of HO2/O-2 Radicals in Aqueous Solution. Journal of Physical and Chemical Reference Data, 14(4), 1041–1100. 10.1063/1.555739

Boldrin, F., Provvedi, R., Cioetto Mazzabò, L., Segafreddo, G., & Manganelli, R. (2020). Tolerance and Persistence to Drugs: A Main Challenge in the Fight Against Mycobacterium Tuberculosis. Frontiers in Microbiology, 11, 1924. 10.3389/fmicb.2020.01924

Bollineni, R. C., Fedorova, M., & Hoffmann, R. (2011). Identification of Carbonylated Peptides by Tandem Mass Spectrometry Using a Precursor Ion-like Scan in Negative Ion Mode. Journal of Proteomics, 74(11), 2351–2359. 10.1016/j.jprot.2011.05.033

Brand, K. A., & Hermfisse, U. (1997). Aerobic Glycolysis by Proliferating Cells: A Protective Strategy against Reactive Oxygen Species. The FASEB Journal, 11(5), 388–395. 10.1096/fasebj.11.5.9141507

Brand, M. D., & Esteves, T. C. (2005). Physiological Functions of the Mitochondrial Uncoupling Proteins UCP2 and UCP3. Cell Metabolism, 2(2), 85–93. 10.1016/j.cmet.2005.06.002

Brandi, G., Sestili, P., Pedrini, M., Salvaggio, L., Cattabeni, F., & Cantoni, O. (1987). The Effect of Temperature or Anoxia on Escherichia Coli Killing Induced by Hydrogen Peroxide. Mutation Research Letters, 190(4), 237–240. 10.1016/0165-7992(87)90002-9

Brauner, A., Fridman, O., Gefen, O., & Balaban, N. Q. (2016). Distinguishing between Resistance, Tolerance and Persistence to Antibiotic Treatment. Nature Reviews Microbiology, 14(5), 320–330. 10.1038/nrmicro.2016.34

Buxton, G. V., Greenstock, C. L., Helman, W. P., & Ross, A. B. (1988). Critical Review of Rate Constants for Reactions of Hydrated Electrons, Hydrogen Atoms and Hydroxyl Radicals (c ·OH/c ·O-in Aqueous Solution. Journal of Physical and Chemical Reference Data, 17(2), 513–886. 10.1063/1.555805

Chance, B., & Williams, G. R. (1956). The Respiratory Chain and Oxidative Phosphorylation. In F. F. Nord (Ed.), Advances in Enzymology and Related Areas of Molecular Biology: Vol. 17. Advances in Enzymology and Related Areas of Molecular Biology (1st ed., pp. 65–134). Wiley. 10.1002/9780470122624.ch2

Chavarría, M., Nikel, P. I., Pérez-Pantoja, D., & De Lorenzo, V. (2013). The Entner–Doudoroff Pathway Empowers Pseudomonas Putida KT2440 with a High Tolerance to Oxidative Stress. Environmental Microbiology, 15(6), 1772–1785. 10.1111/1462-2920.12069

Cheignon, C., Tomas, M., Bonnefont-Rousselot, D., Faller, P., Hureau, C., & Collin, F. (2018). Oxidative Stress and the Amyloid Beta Peptide in Alzheimer’s Disease. Redox Biology, 14, 450–464. 10.1016/j.redox.2017.10.014

Chen, H.-Y. (2019). Why the Reactive Oxygen Species of the Fenton Reaction Switches from Oxoiron(IV) Species to Hydroxyl Radical in Phosphate Buffer Solutions? A Computational Rationale. ACS Omega, 4(9), 14105–14113. 10.1021/acsomega.9b02023

Christodoulou, D., Link, H., Fuhrer, T., Kochanowski, K., Gerosa, L., & Sauer, U. (2018). Reserve Flux Capacity in the Pentose Phosphate Pathway Enables Escherichia Coli’s Rapid Response to Oxidative Stress. Cell Systems, 6(5), 569–578.e7. 10.1016/j.cels.2018.04.009

Cioffi, F., Adam, R. H. I., & Broersen, K. (2019). Molecular Mechanisms and Genetics of Oxidative Stress in Alzheimer’s Disease. Journal of Alzheimer’s Disease, 72(4), 981–1017. 10.3233/JAD-190863

Collin, F. (2019). Chemical Basis of Reactive Oxygen Species Reactivity and Involvement in Neurodegenerative Diseases. International Journal of Molecular Sciences, 20(10), 2407. 10.3390/ijms20102407

Dahlgren, C., Karlsson, A., & Bylund, J. (2019). Intracellular Neutrophil Oxidants: From Laboratory Curiosity to Clinical Reality. The Journal of Immunology, 202(11), 3127–3134. 10.4049/jimmunol.1900235

Davalli, P., Marverti, G., Lauriola, A., & D’Arca, D. (2018). Targeting Oxidatively Induced DNA Damage Response in Cancer: Opportunities for Novel Cancer Therapies. Oxidative Medicine and Cellular Longevity, 2018(1), 2389523. 10.1155/2018/2389523

DeBerardinis, R. J., & Chandel, N. S. (2020). We Need to Talk about the Warburg Effect. Nature Metabolism, 2(2), 127–129. 10.1038/s42255-020-0172-2

Desouki, A. A., Jarre, F., Gelius-Dietrich, G., & Lercher, M. J. (2015). CycleFreeFlux: Efficient Removal of Thermodynamically Infeasible Loops from Flux Distributions. Bioinformatics, 31(13), 2159–2165. 10.1093/bioinformatics/btv096

Diaz-Ruiz, R., Rigoulet, M., & Devin, A. (2011). The Warburg and Crabtree Effects: On the Origin of Cancer Cell Energy Metabolism and of Yeast Glucose Repression. Biochimica Et Biophysica Acta (BBA) - Bioenergetics, 1807(6), 568–576. 10.1016/j.bbabio.2010.08.010

Diaz-Vegas, A., Sanchez-Aguilera, P., Krycer, J. R., Morales, P. E., Monsalves-Alvarez, M., Cifuentes, M., Rothermel, B. A., & Lavandero, S. (2020). Is Mitochondrial Dysfunction a Common Root of Noncommunicable Chronic Diseases?. Endocrine Reviews, 41(3), bnaa5. 10.1210/endrev/bnaa005

Fanous, J., Claudi, B., Tripathi, V., Li, J., Goormaghtigh, F., & Bumann, D. (2025). Limited Impact of Salmonella Stress and Persisters on Antibiotic Clearance. Nature, 639(8053), 181–189. 10.1038/s41586-024-08506-6

Fasnacht, M., & Polacek, N. (2021). Oxidative Stress in Bacteria and the Central Dogma of Molecular Biology. Frontiers in Molecular Biosciences, 8, 671037. 10.3389/fmolb.2021.671037

Fenton, H. J. H. (1894). LXXIII.—Oxidation of Tartaric Acid in Presence of Iron. J. Chem. Soc., Trans., 65(0), 899–910. 10.1039/CT8946500899

Flamholz, A. I., Dugan, E., Blikstad, C., Gleizer, S., Ben-Nissan, R., Amram, S., Antonovsky, N., Ravishankar, S., Noor, E., Bar-Even, A., Milo, R., & Savage, D. F. (2020). Functional Reconstitution of a Bacterial CO2 Concentrating Mechanism in Escherichia Coli. Elife, 9, e59882. 10.7554/eLife.59882

Forenzo, C., & Larsen, J. (2024). Bridging Clinical Radiotherapy and Space Radiation Therapeutics through Reactive Oxygen Species (ROS)-Triggered Delivery. Free Radical Biology and Medicine, 219, 88–103. 10.1016/j.freeradbiomed.2024.04.219

Franceschi, C., Garagnani, P., Morsiani, C., Conte, M., Santoro, A., Grignolio, A., Monti, D., Capri, M., & Salvioli,S. (2018). The Continuum of Aging and Age-Related Diseases: Common Mechanisms but Different Rates. Frontiers in Medicine, 5, 61. 10.3389/fmed.2018.00061

Garrison, W. M. (1987). Reaction Mechanisms in the Radiolysis of Peptides, Polypeptides, and Proteins. Chemical Reviews, 87(2), 381–398. 10.1021/cr00078a006

Ge, H., Wu, P., Qian, H., & Xie, X. S. (2018). Relatively Slow Stochastic Gene-State Switching in the Presence of Positive Feedback Significantly Broadens the Region of Bimodality through Stabilizing the Uninduced Phenotypic State. PLOS Computational Biology, 14(3), e1006051. 10.1371/journal.pcbi.1006051

Geary, L. E., & Meister, A. (1977). On the Mechanism of Glutamine-Dependent Reductive Amination of Alpha-Ketoglutarate Catalyzed by Glutamate Synthase. Journal of Biological Chemistry, 252(10), 3501–3508. 10.1016/S0021-9258(17)40419-4

Gefen, O., & Balaban, N. Q. (2009). The Importance of Being Persistent: Heterogeneity of Bacterial Populations under Antibiotic Stress. FEMS Microbiology Reviews, 33(4), 704–717. 10.1111/j.1574-6976.2008.00156.x

Gianazza, E., Brioschi, M., Martinez Fernandez, A., Casalnuovo, F., Altomare, A., Aldini, G., & Banfi, C. (2021). Lipid Peroxidation in Atherosclerotic Cardiovascular Diseases. Antioxidants & Redox Signaling, 34(1), 49–98. 10.1089/ars.2019.7955

Goelzer, A., Muntel, J., Chubukov, V., Jules, M., Prestel, E., Nölker, R., Mariadassou, M., Aymerich, S., Hecker, M., Noirot, P., Becher, D., & Fromion, V. (2015). Quantitative Prediction of Genome-Wide Resource Allocation in Bacteria. Metabolic Engineering, 32, 232–243. 10.1016/j.ymben.2015.10.003

Goormaghtigh, F., & Van Melderen, L. (2019). Single-Cell Imaging and Characterization of Escherichia Coli Persister Cells to Ofloxacin in Exponential Cultures. Science Advances, 5(6), eaav9462. 10.1126/sciadv.aav9462

Gray, B., & Carmichael, A. J. (1992). Kinetics of Superoxide Scavenging by Dismutase Enzymes and Manganese Mimics Determined by Electron Spin Resonance. Biochemical Journal, 281(3), 795–802. 10.1042/bj2810795

Green, R., Wang, H., Botchey, C., Zhang, S. N. N., Wadsworth, C., Tyrrell, F., Letton, J., McBain, A. J., Paszek, P., Krašovec, R., & Knight, C. G. (2024). Collective Peroxide Detoxification Determines Microbial Mutation Rate Plasticity in E. Coli. PLOS Biology, 22(7), e3002711. 10.1371/journal.pbio.3002711

Guo, K., Wu, S., Qin, W., Xie, R., Wu, Y., Li, X., Ouyang, G., & Fang, J. (2024). Overlooked Generation of Reactive Oxidative Species from Water and Dioxygen by Far UV Light. Environmental Science & Technology, 58(50), 22431–22441. 10.1021/acs.est.4c06404

Haque, M. F., Tarusawa, T., Ushida, C., Ito, S., & Himeno, H. (2025). cAMP-CRP-Activated E. Coli Causes Growth Arrest under Stress Conditions. Frontiers in Microbiology, 16, 1597530. 10.3389/fmicb.2025.1597530

Harcombe, W., Riehl, W., Dukovski, I., Granger, B., Betts, A., Lang, A., Bonilla, G., Kar, A., Leiby, N., Mehta, P., Marx, C., & Segrè, D. (2014). Metabolic Resource Allocation in Individual Microbes Determines Ecosystem Interactions and Spatial Dynamics. Cell Reports, 7(4), 1104–1115. 10.1016/j.celrep.2014.03.070

Hashimoto, M., Ichimura, T., Mizoguchi, H., Tanaka, K., Fujimitsu, K., Keyamura, K., Ote, T., Yamakawa, T., Yamazaki, Y., Mori, H., Katayama, T., & Kato, J.-i. (2005). Cell Size and Nucleoid Organization of Engineered Escherichia Coli Cells with a Reduced Genome. Molecular Microbiology, 55(1), 137–149. 10.1111/j.1365-2958.2004.04386.x

Hawkins, C. L., & Davies, M. J. (2019). Detection, Identification, and Quantification of Oxidative Protein Modifications. Journal of Biological Chemistry, 294(51), 19683–19708. 10.1074/jbc.REV119.006217

Heirendt, L., Arreckx, S., Pfau, T., Mendoza, S. N., Richelle, A., Heinken, A., Haraldsdóttir, H. S., Wachowiak, J., Keating, S. M., Vlasov, V., Magnusdóttir, S., Ng, C. Y., Preciat, G., Žagare, A., Chan, S. H. J., Aurich, M. K., Clancy, C. M., Modamio, J., Sauls, J. T.,…Fleming, R. M. T. (2019). Creation and Analysis of Biochemical Constraint-Based Models Using the COBRA Toolbox v.3.0. Nature Protocols, 14(3), 639–702. 10.1038/s41596-018-0098-2

Höhn, A., Tramutola, A., & Cascella, R. (2020). Proteostasis Failure in Neurodegenerative Diseases: Focus on Oxidative Stress. Oxidative Medicine and Cellular Longevity, 2020, 1–21. 10.1155/2020/5497046

Imlay, J. A. (1995). A Metabolic Enzyme That Rapidly Produces Superoxide, Fumarate Reductase of Escherichia Coli. Journal of Biological Chemistry, 270(34), 19767–19777. 10.1016/S0021-9258(18)94548-5

Imlay, J. A. (2006). Iron-sulphur Clusters and the Problem with Oxygen. Molecular Microbiology, 59(4), 1073–1082. 10.1111/j.1365-2958.2006.05028.x

Imlay, J. A. (2008). Cellular Defenses against Superoxide and Hydrogen Peroxide. Annual Review of Biochemistry, 77(1), 755–776. 10.1146/annurev.biochem.77.061606.161055

Imlay, J. A. (2013). The Molecular Mechanisms and Physiological Consequences of Oxidative Stress: Lessons from a Model Bacterium. Nature Reviews Microbiology, 11(7), 443–454. 10.1038/nrmicro3032

Imlay, J. A. (2019). Where in the World Do Bacteria Experience Oxidative Stress?. Environmental Microbiology, 21(2), 521–530. 10.1111/1462-2920.14445

Imlay, J. A., & Linn, S. (1986). Bimodal Pattern of Killing of DNA-Repair-Defective or Anoxically Grown Escherichia Coli by Hydrogen Peroxide. Journal of Bacteriology, 166(2), 519–527. 10.1128/jb.166.2.519-527.1986

Ito, F., Sono, Y., & Ito, T. (2019). Measurement and Clinical Significance of Lipid Peroxidation as a Biomarker of Oxidative Stress: Oxidative Stress in Diabetes, Atherosclerosis, and Chronic Inflammation. Antioxidants, 8(3), 72. 10.3390/antiox8030072

Jastroch, M., Divakaruni, A. S., Mookerjee, S., Treberg, J. R., & Brand, M. D. (2010). Mitochondrial Proton and Electron Leaks. Essays in Biochemistry, 47, 53–67. 10.1042/bse0470053

Jiang, B., & Moskovitz, J. (2018). The Functions of the Mammalian Methionine Sulfoxide Reductase System and Related Diseases. Antioxidants, 7(9), 122. 10.3390/antiox7090122

Jozefczuk, S., Klie, S., Catchpole, G., Szymanski, J., Cuadros-Inostroza, A., Steinhauser, D., Selbig, J., & Willmitzer,L. (2010). Metabolomic and Transcriptomic Stress Response of Escherichia Coli. Molecular Systems Biology, 6(1), MSB201018. 10.1038/msb.2010.18

Juan, C. A., Pérez De La Lastra, J. M., Plou, F. J., & Pérez-Lebeña, E. (2021). The Chemistry of Reactive Oxygen Species (ROS) Revisited: Outlining Their Role in Biological Macromolecules (DNA, Lipids and Proteins) and Induced Pathologies. International Journal of Molecular Sciences, 22(9), 4642. 10.3390/ijms22094642

Kaneko, M., Kodama, M., & Inoue, F. (1994). Bimodal Pattern of Killing of Chinese Hamster V79 Variant Cells by Hydrogen Peroxide. Free Radical Research, 20(4), 229–239. 10.3109/10715769409147519

Kao, Y.-C., Ho, P.-C., Tu, Y.-K., Jou, I.-M., & Tsai, K.-J. (2020). Lipids and Alzheimer’s Disease. International Journal of Molecular Sciences, 21(4), 1505. 10.3390/ijms21041505

Keaty, T. C., & Jensen, P. A. (2020). Gapsplit: Efficient Random Sampling for Non-Convex Constraint-Based Models. Bioinformatics, 36(8), 2623–2625. 10.1093/bioinformatics/btz971

Kehrer, J. P. (2000). The Haber–Weiss Reaction and Mechanisms of Toxicity. Toxicology, 149(1), 43–50. 10.1016/S0300-483X(00)00231-6

Koppenol, W. (2001). The Haber-Weiss Cycle – 70 Years Later. Redox Report, 6(4), 229–234. 10.1179/135100001101536373

Korshunov, S., & Imlay, J. A. (2010). Two Sources of Endogenous Hydrogen Peroxide in Escherichia Coli. Molecular Microbiology, 75(6), 1389–1401. 10.1111/j.1365-2958.2010.07059.x

Krumova, K., & Cosa, G. (2016). Chapter 1. Overview of Reactive Oxygen Species. In S. Nonell & C. Flors (Eds.), Singlet Oxygen: Vol. 1. Singlet Oxygen (pp. 1–21). Royal Society of Chemistry. 10.1039/9781782622208-00001

Kumar, S. R., & Imlay, J. A. (2013). How Escherichia Coli Tolerates Profuse Hydrogen Peroxide Formation by a Catabolic Pathway. Journal of Bacteriology, 195(20), 4569–4579. 10.1128/JB.00737-13

Kussell, E., & Leibler, S. (2005). Phenotypic Diversity, Population Growth, and Information in Fluctuating Environments. Science, 309(5743), 2075–2078. 10.1126/science.1114383

Kussmaul, L., & Hirst, J. (2006). The Mechanism of Superoxide Production by NADH:Ubiquinone Oxidoreductase (Complex I) from Bovine Heart Mitochondria. Proceedings of the National Academy of Sciences, 103(20), 7607–7612. 10.1073/pnas.0510977103

Lam, P.-L., Wong, R.-M., Lam, K.-H., Hung, L.-K., Wong, M.-M., Yung, L.-H., Ho, Y.-W., Wong, W.-Y., Hau, D.-P., Gambari, R., & Chui, C.-H. (2020). The Role of Reactive Oxygen Species in the Biological Activity of Antimicrobial Agents: An Updated Mini Review. Chemico-Biological Interactions, 320, 109023. 10.1016/j.cbi.2020.109023

Lawless, C., Greaves, L., Reeve, A. K., Turnbull, D. M., & Vincent, A. E. (2020). The Rise and Rise of Mitochondrial DNA Mutations. Open Biology, 10(5), 200061. 10.1098/rsob.200061

Lee, P., Vousden, K. H., & Cheung, E. C. (2014). TIGAR, TIGAR, Burning Bright. Cancer & Metabolism, 2(1), 1. 10.1186/2049-3002-2-1

Li, X., & Imlay, J. A. (2018). Improved Measurements of Scant Hydrogen Peroxide Enable Experiments That Define Its Threshold of Toxicity for Escherichia Coli. Free Radical Biology and Medicine, 120, 217–227. 10.1016/j.freeradbiomed.2018.03.025

Lin, T.-Y. (1999). G33D Mutant Thioredoxin Primarily Affects the Kinetics of Reaction with Thioredoxin Reductase. Probing the Structure of the Mutant Protein. Biochemistry, 38(47), 15508–15513. 10.1021/bi9907678

Linley, E., Denyer, S. P., McDonnell, G., Simons, C., & Maillard, J.-Y. (2012). Use of Hydrogen Peroxide as a Biocide: New Consideration of Its Mechanisms of Biocidal Action. Journal of Antimicrobial Chemotherapy, 67(7), 1589–1596. 10.1093/jac/dks129

Liochev, S. I., & Fridovich, I. (2002). The Haber-Weiss Cycle—70 Years Later: An Alternative View. Redox Report, 7(1), 55–57. 10.1179/135100002125000190

Liu, X., Ramsey, M. M., Chen, X., Koley, D., Whiteley, M., & Bard, A. J. (2011). Real-Time Mapping of a Hydrogen Peroxide Concentration Profile across a Polymicrobial Bacterial Biofilm Using Scanning Electrochemical Microscopy. Proceedings of the National Academy of Sciences, 108(7), 2668–2673. 10.1073/pnas.1018391108

Loewen, P. C., Switala, J., & Triggs-Raine, B. L. (1985). Catalases HPI and HPII in Escherichia Coli Are Induced Independently. Archives of Biochemistry and Biophysics, 243(1), 144–149. 10.1016/0003-9861(85)90782-9

Lu, Z., & Imlay, J. A. (2021). When Anaerobes Encounter Oxygen: Mechanisms of Oxygen Toxicity, Tolerance and Defence. Nature Reviews Microbiology, 19(12), 774–785. 10.1038/s41579-021-00583-y

M9 Minimal Medium (Standard). (2010). Cold Spring Harbor Protocols, 2010(8), pdb.rec12295. 10.1101/pdb.rec12295

Magnani, F., & Mattevi, A. (2019). Structure and Mechanisms of ROS Generation by NADPH Oxidases. Current Opinion in Structural Biology, 59, 91–97. 10.1016/j.sbi.2019.03.001

Mahadevan, R., & Schilling, C. (2003). The Effects of Alternate Optimal Solutions in Constraint-Based Genome-Scale Metabolic Models. Metabolic Engineering, 5(4), 264–276. 10.1016/j.ymben.2003.09.002

Mailloux, R. J. (2020). An Update on Mitochondrial Reactive Oxygen Species Production. Antioxidants, 9(6), 472. 10.3390/antiox9060472

Maisonneuve, E., Ducret, A., Khoueiry, P., Lignon, S., Longhi, S., Talla, E., & Dukan, S. (2009). Rules Governing Selective Protein Carbonylation. Plos ONE, 4(10), e7269. 10.1371/journal.pone.0007269

Marinos, G., Kaleta, C., & Waschina, S. (2020). Defining the Nutritional Input for Genome-Scale Metabolic Models: A Roadmap. PLOS ONE, 15(8), e236890. 10.1371/journal.pone.0236890

Massey, V., Strickland, S., Mayhew, S., Howell, L., Engel, P., Matthews, R., Schuman, M., & Sullivan, P. (1969). The Production of Superoxide Anion Radicals in the Reaction of Reduced Flavins and Flavoproteins with Molecular Oxygen. Biochemical and Biophysical Research Communications, 36(6), 891–897.10.1016/0006-291X(69)90287-3

Matallana-Surget, S., Cavicchioli, R., Fauconnier, C., Wattiez, R., Leroy, B., Joux, F., Raftery, M. J., & Lebaron, P. (2013). Shotgun Redox Proteomics: Identification and Quantitation of Carbonylated Proteins in the UVB-Resistant Marine Bacterium, Photobacterium Angustum S14. Plos ONE, 8(7), e68112. 10.1371/journal.pone.0068112

Matilla, M. A., & Krell, T. (2018). The Effect of Bacterial Chemotaxis on Host Infection and Pathogenicity. FEMS Microbiology Reviews, 42(1). 10.1093/femsre/fux052

Messner, K. R., & Imlay, J. A. (1999). The Identification of Primary Sites of Superoxide and Hydrogen Peroxide Formation in the Aerobic Respiratory Chain and Sulfite Reductase Complex of Escherichia Coli. Journal of Biological Chemistry, 274(15), 10119–10128. 10.1074/jbc.274.15.10119

Messner, K. R., & Imlay, J. A. (2002). Mechanism of Superoxide and Hydrogen Peroxide Formation by Fumarate Reductase, Succinate Dehydrogenase, and Aspartate Oxidase. Journal of Biological Chemistry, 277(45), 42563–42571. 10.1074/jbc.M204958200

Milkovic, L., Cipak Gasparovic, A., Cindric, M., Mouthuy, P.-A., & Zarkovic, N. (2019). Short Overview of ROS as Cell Function Regulators and Their Implications in Therapy Concepts. Cells, 8(8), 793. 10.3390/cells8080793

Mirzaei, H., & Regnier, F. (2006). Creation of Allotypic Active Sites during Oxidative Stress. Journal of Proteome Research, 5(9), 2159–2168. 10.1021/pr060021d

Mishra, S., & Imlay, J. (2012). Why Do Bacteria Use so Many Enzymes to Scavenge Hydrogen Peroxide?. Archives of Biochemistry and Biophysics, 525(2), 145–160. 10.1016/j.abb.2012.04.014

Mizrachi, A., Graff Van Creveld, S., Shapiro, O. H., Rosenwasser, S., & Vardi, A. (2019). Light-Dependent Single-Cell Heterogeneity in the Chloroplast Redox State Regulates Cell Fate in a Marine Diatom. Elife, 8, e47732. 10.7554/eLife.47732

Molenaar, D., Van Berlo, R., De Ridder, D., & Teusink, B. (2009). Shifts in Growth Strategies Reflect Tradeoffs in Cellular Economics. Molecular Systems Biology, 5(1), MSB200982. 10.1038/msb.2009.82

Monk, J. M., Lloyd, C. J., Brunk, E., Mih, N., Sastry, A., King, Z., Takeuchi, R., Nomura, W., Zhang, Z., Mori, H., Feist, A. M., & Palsson, B. O. (2017). iML1515, a Knowledgebase That Computes Escherichia Coli Traits. Nature Biotechnology, 35(10), 904–908. 10.1038/nbt.3956

Moskovitz, J. (2005). Methionine Sulfoxide Reductases: Ubiquitous Enzymes Involved in Antioxidant Defense, Protein Regulation, and Prevention of Aging-Associated Diseases. Biochimica Et Biophysica Acta (BBA) - Proteins and Proteomics, 1703(2), 213–219. 10.1016/j.bbapap.2004.09.003

Moskovitz, J., Rahman, M. A., Strassman, J., Yancey, S. O., Kushner, S. R., Brot, N., & Weissbach, H. (1995). Escherichia Coli Peptide Methionine Sulfoxide Reductase Gene: Regulation of Expression and Role in Protecting against Oxidative Damage. Journal of Bacteriology, 177(3), 502–507. 10.1128/jb.177.3.502-507.1995

Neidhardt, F. C., Ingraham, J. L., & Schaechter, M. (1990). Physiology of the Bacterial Cell: A Molecular Approach. Sinauer Associates.

Nesci, S., & Rubattu, S. (2024). UCP2, a Member of the Mitochondrial Uncoupling Proteins: An Overview from Physiological to Pathological Roles. Biomedicines, 12(6), 1307. 10.3390/biomedicines12061307

Nguyen, G. T., Green, E. R., & Mecsas, J. (2017). Neutrophils to the ROScue: Mechanisms of NADPH Oxidase Activation and Bacterial Resistance. Frontiers in Cellular and Infection Microbiology, 7, 373. 10.3389/fcimb.2017.00373

Nissanka, N., & Moraes, C. T. (2018). Mitochondrial DNA Damage and Reactive Oxygen Species in Neurode-generative Disease!. FEBS Letters, 592(5), 728–742. 10.1002/1873-3468.12956

Orth, J. D., Conrad, T. M., Na, J., Lerman, J. A., Nam, H., Feist, A. M., & Palsson, B. Ø. (2011). A Comprehensive Genome-scale Reconstruction of Escherichia Coli Metabolism—2011. Molecular Systems Biology, 7(1), MSB201165. 10.1038/msb.2011.65

Orth, J. D., Thiele, I., & Palsson, B. Ø. (2010). What Is Flux Balance Analysis?. Nature Biotechnology, 28(3), 245–248. 10.1038/nbt.1614

Oyarzún, D. A., & Chaves, M. (2015). Design of a Bistable Switch to Control Cellular Uptake. Journal of the Royal Society Interface, 12(113), 20150618. 10.1098/rsif.2015.0618

Palma, F. R., He, C., Danes, J. M., Paviani, V., Coelho, D. R., Gantner, B. N., & Bonini, M. G. (2020). Mitochondrial Superoxide Dismutase: What the Established, the Intriguing, and the Novel Reveal About a Key Cellular Redox Switch. Antioxidants & Redox Signaling, 32(10), 701–714. 10.1089/ars.2019.7962

Perez-Samper, G., Cerulus, B., Jariani, A., Vermeersch, L., Barrajón Simancas, N., Bisschops, M. M. M., Van Den Brink, J., Solis-Escalante, D., Gallone, B., De Maeyer, D., Van Bael, E., Wenseleers, T., Michiels, J., Marchal, K., Daran-Lapujade, P., & Verstrepen, K. J. (2018). The Crabtree Effect Shapes the Saccharomyces Cerevisiae Lag Phase during the Switch between Different Carbon Sources. Mbio, 9(5), e1331–18. 10.1128/mBio.01331-18

Perillo, B., Di Donato, M., Pezone, A., Di Zazzo, E., Giovannelli, P., Galasso, G., Castoria, G., & Migliaccio, A. (2020). ROS in Cancer Therapy: The Bright Side of the Moon. Experimental & Molecular Medicine, 52(2), 192–203. 10.1038/s12276-020-0384-2

Poetsch, A. R. (2020). The Genomics of Oxidative DNA Damage, Repair, and Resulting Mutagenesis. Computational and Structural Biotechnology Journal, 18, 207–219. 10.1016/j.csbj.2019.12.013

Quigley, J., & Lewis, K. (2022). Noise in a Metabolic Pathway Leads to Persister Formation in Mycobacterium Tuberculosis. Microbiology Spectrum, 10(5), e2948–22. 10.1128/spectrum.02948-22

Radzikowski, J. L., Schramke, H., & Heinemann, M. (2017). Bacterial Persistence from a System-Level Perspective. Current Opinion in Biotechnology, 46, 98–105. 10.1016/j.copbio.2017.02.012

Radzikowski, J. L., Vedelaar, S., Siegel, D., Ortega, Á. D., Schmidt, A., & Heinemann, M. (2016). Bacterial Persistence Is an Active Sigma-S Stress Response to Metabolic Flux Limitation. Molecular Systems Biology, 12(9), MSB166998. 10.15252/msb.20166998

Raser, J. M., & O’Shea, E. K. (2005). Noise in Gene Expression: Origins, Consequences, and Control. Science, 309(5743), 2010–2013. 10.1126/science.1105891

Read, A. D., Bentley, R. E., Archer, S. L., & Dunham-Snary, K. J. (2021). Mitochondrial Iron–Sulfur Clusters: Structure, Function, and an Emerging Role in Vascular Biology. Redox Biology, 47, 102164.10.1016/j.redox.2021.102164

Rodríguez-Rojas, A., Kim, J. J., Johnston, P. R., Makarova, O., Eravci, M., Weise, C., Hengge, R., & Rolff, J. (2020). Non-Lethal Exposure to H2O2 Boosts Bacterial Survival and Evolvability against Oxidative Stress. PLOS Genetics, 16(3), e1008649. 10.1371/journal.pgen.1008649

Rotem, E., Loinger, A., Ronin, I., Levin-Reisman, I., Gabay, C., Shoresh, N., Biham, O., & Balaban, N. Q. (2010). Regulation of Phenotypic Variability by a Threshold-Based Mechanism Underlies Bacterial Persistence. Proceedings of the National Academy of Sciences, 107(28), 12541–12546. 10.1073/pnas.1004333107

Roth, M., Goodall, E. C. A., Pullela, K., Jaquet, V., François, P., Henderson, I. R., & Krause, K.-H. (2022). Transposon-Directed Insertion-Site Sequencing Reveals Glycolysis Gene gpmA as Part of the H2O2 Defense Mechanisms in Escherichia Coli. Antioxidants, 11(10), 2053. 10.3390/antiox11102053

Sahu, S. K., & Behuria, H. G. (2018). Biochemical Characterization of H2O2-Induced Oxidative Stress in E.Coli. Journal of Applied Microbiology and Biochemistry, 2(3). 10.21767/2576-1412.100026

Salehi, F., Behboudi, H., Kavoosi, G., & Ardestani, S. K. (2018). Oxidative DNA Damage Induced by ROS-Modulating Agents with the Ability to Target DNA: A Comparison of the Biological Characteristics of Citrus Pectin and Apple Pectin. Scientific Reports, 8(1), 13902. 10.1038/s41598-018-32308-2

Santos-Sánchez, N. F., Salas-Coronado, R., Villanueva-Cañongo, C., & Hernández-Carlos, B. (2019). Antioxidant Compounds and Their Antioxidant Mechanism. In E. Shalaby (Ed.), Antioxidants: Antioxidants. IntechOpen. 10.5772/intechopen.85270

Schittenhelm, D., Neuss-Radu, M., Verma, N., Pink, M., & Schmitz-Spanke, S. (2019). ROS and Pentose Phosphate Pathway: Mathematical Modelling of the Metabolic Regulation in Response to Xenobiotic-Induced Oxidative Stress and the Proposed Impact of the Gluconate Shunt. Free Radical Research, 53(9–10), 979–992. 10.1080/10715762.2019.1660777

Schleicher, J. (2021). Introduction to In Silico Modeling to Study ROS Dynamics. In J. Espada (Ed.), Reactive Oxygen Species: Vol. 2202. Reactive Oxygen Species (pp. 1–32). Springer US. 10.1007/978-1-0716-0896-8_1

Schuetz, R., Zamboni, N., Zampieri, M., Heinemann, M., & Sauer, U. (2012). Multidimensional Optimality of Microbial Metabolism. Science, 336(6081), 601–604. 10.1126/science.1216882

Seaver, L. C., & Imlay, J. A. (2001b). Alkyl Hydroperoxide Reductase Is the Primary Scavenger of Endogenous Hydrogen Peroxide in Escherichia Coli. Journal of Bacteriology, 183(24), 7173–7181. 10.1128/JB.183.24.7173-7181.2001

Seaver, L. C., & Imlay, J. A. (2001a). Hydrogen Peroxide Fluxes and Compartmentalization inside Growing Escherichia Coli. Journal of Bacteriology, 183(24), 7182–7189. 10.1128/JB.183.24.7182-7189.2001

Segal, A. W. (2008). The Function of the NADPH Oxidase of Phagocytes and Its Relationship to Other NOXs in Plants, Invertebrates, and Mammals. The International Journal of Biochemistry & Cell Biology, 40(4), 604–618. 10.1016/j.biocel.2007.10.003

Sezonov, G., Joseleau-Petit, D., & D’Ari, R. (2007). Escherichia Coli Physiology in Luria-Bertani Broth. Journal of Bacteriology, 189(23), 8746–8749. 10.1128/JB.01368-07

Shabestary, K., Klemm, C., Carling, B., Marshall, J., Savigny, J., Storch, M., & Ledesma-Amaro, R. (2024). Phenotypic Heterogeneity Follows a Growth-Viability Tradeoff in Response to Amino Acid Identity. Nature Communications, 15(1), 6515. 10.1038/s41467-024-50602-8

Sharma, P., & Sampath, H. (2019). Mitochondrial DNA Integrity: Role in Health and Disease. Cells, 8(2), 100. 10.3390/cells8020100

Shiratori, R., Furuichi, K., Yamaguchi, M., Miyazaki, N., Aoki, H., Chibana, H., Ito, K., & Aoki, S. (2019). Glycolytic Suppression Dramatically Changes the Intracellular Metabolic Profile of Multiple Cancer Cell Lines in a Mitochondrial Metabolism-Dependent Manner. Scientific Reports, 9(1), 18699. 10.1038/s41598-019-55296-3

Shultis, M. W., Mulholland, C. V., & Berney, M. (2022). Are All Antibiotic Persisters Created Equal?. Frontiers in Cellular and Infection Microbiology, 12, 933458. 10.3389/fcimb.2022.933458

Sies, H., Mailloux, R. J., & Jakob, U. (2024). Fundamentals of Redox Regulation in Biology. Nature Reviews Molecular Cell Biology, 25(9), 701–719. 10.1038/s41580-024-00730-2

Snezhkina, A. V., Kudryavtseva, A. V., Kardymon, O. L., Savvateeva, M. V., Melnikova, N. V., Krasnov, G. S., & Dmitriev, A. A. (2019). ROS Generation and Antioxidant Defense Systems in Normal and Malignant Cells. Oxidative Medicine and Cellular Longevity, 2019, 1–17. 10.1155/2019/6175804

Sridhar, S., Bhalla, P., Kullu, J., Veerapaneni, S., Sahoo, S., Bhatt, N., & Suraishkumar, G. (2023). A Reactive Species Reactions Module for Integration into Genome-Scale Metabolic Models for Improved Insights: Application to Cancer. Metabolic Engineering, 80, 78–93. 10.1016/j.ymben.2023.08.006

Srinivas, U. S., Tan, B. W., Vellayappan, B. A., & Jeyasekharan, A. D. (2019). ROS and the DNA Damage Response in Cancer. Redox Biology, 25, 101084. 10.1016/j.redox.2018.101084

Stadtman, E. R., & Levine, R. L. (2003). Free Radical-Mediated Oxidation of Free Amino Acids and Amino Acid Residues in Proteins. Amino Acids, 25(3–4), 207–218. 10.1007/s00726-003-0011-2

Szabó, E., & Ambrus, A. (2023). Lipoamide Dehydrogenase (LADH) Deficiency: Medical Perspectives of the Structural and Functional Characterization of LADH and Its Pathogenic Variants. Biologia Futura, 74(1–2), 109–118. 10.1007/s42977-023-00155-6

Taborsky, G., & McCollum, K. (1973). Oxidative Modification of Proteins in the Presence of Ferrous Ion and Air. Effect of Ionic Constituents of the Reaction Medium on the Nature of the Oxidation Products. Biochemistry, 12(7), 1341–1348. 10.1021/bi00731a014

Takhaveev, V., & Heinemann, M. (2018). Metabolic Heterogeneity in Clonal Microbial Populations. Current Opinion in Microbiology, 45, 30–38. 10.1016/j.mib.2018.02.004

Tarrago, L., Kaya, A., Weerapana, E., Marino, S. M., & Gladyshev, V. N. (2012). Methionine Sulfoxide Reductases Preferentially Reduce Unfolded Oxidized Proteins and Protect Cells from Oxidative Protein Unfolding. Journal of Biological Chemistry, 287(29), 24448–24459. 10.1074/jbc.M112.374520

Taverne, Y. J. H. J., Bogers, A. J. J. C., Duncker, D. J., & Merkus, D. (2013). Reactive Oxygen Species and the Cardiovascular System. Oxidative Medicine and Cellular Longevity, 2013, 1–15. 10.1155/2013/862423

Temple, A., Yen, T. Y., & Gronert, S. (2006). Identification of Specific Protein Carbonylation Sites in Model Oxidations of Human Serum Albumin. Journal of the American Society for Mass Spectrometry, 17(8), 1172–1180. 10.1016/j.jasms.2006.04.030

Theurey, P., & Pizzo, P. (2018). The Aging Mitochondria. Genes, 9(1), 22. 10.3390/genes9010022

To, T.-L., & Maheshri, N. (2010). Noise Can Induce Bimodality in Positive Transcriptional Feedback Loops Without Bistability. Science, 327(5969), 1142–1145. 10.1126/science.1178962

Tong, H., Chen, W., Merritt, J., Qi, F., Shi, W., & Dong, X. (2007). Streptococcus Oligofermentans Inhibits Streptococcus Mutans through Conversion of Lactic Acid into Inhibitory H2O2: A Possible Counteroffensive Strategy for Interspecies Competition. Molecular Microbiology, 63(3), 872–880. 10.1111/j.1365-2958.2006.05546.x

Tonn, M. K., Thomas, P., Barahona, M., & Oyarzún, D. A. (2019). Stochastic Modelling Reveals Mechanisms of Metabolic Heterogeneity. Communications Biology, 2(1), 108. 10.1038/s42003-019-0347-0

Turrens, J. F. (2003). Mitochondrial Formation of Reactive Oxygen Species. The Journal of Physiology, 552(2), 335–344. 10.1113/jphysiol.2003.049478

Turrens, J. F., Alexandre, A., & Lehninger, A. L. (1985). Ubisemiquinone Is the Electron Donor for Superoxide Formation by Complex III of Heart Mitochondria. Archives of Biochemistry and Biophysics, 237(2), 408–414. 10.1016/0003-9861(85)90293-0

Uhl, L., & Dukan, S. (2016). Hydrogen Peroxide Induced Cell Death: The Major Defences Relative Roles and Consequences in E. Coli. PLOS ONE, 11(8), e159706. 10.1371/journal.pone.0159706

Uhl, L., Gerstel, A., Chabalier, M., & Dukan, S. (2015). Hydrogen Peroxide Induced Cell Death: One or Two Modes of Action?. Heliyon, 1(4), e49. 10.1016/j.heliyon.2015.e00049

Vallejo, F. A., Vanni, S., & Graham, R. M. (2021). UCP2 as a Potential Biomarker for Adjunctive Metabolic Therapies in Tumor Management. Frontiers in Oncology, 11, 640720. 10.3389/fonc.2021.640720

Veening, J.-W., Stewart, E. J., Berngruber, T. W., Taddei, F., Kuipers, O. P., & Hamoen, L. W. (2008). Bet-Hedging and Epigenetic Inheritance in Bacterial Cell Development. Proceedings of the National Academy of Sciences, 105(11), 4393–4398. 10.1073/pnas.0700463105

Vielma, J. P. (2015). Mixed Integer Linear Programming Formulation Techniques. SIAM Review, 57(1), 3–57. 10.1137/130915303

Wang, H., Jiang, H., Van De Gucht, M., & De Ridder, M. (2019). Hypoxic Radioresistance: Can ROS Be the Key to Overcome It?. Cancers, 11(1), 112. 10.3390/cancers11010112

Wehrens, M., Büke, F., Nghe, P., & Tans, S. J. (2018). Stochasticity in Cellular Metabolism and Growth: Approaches and Consequences. Current Opinion in Systems Biology, 8, 131–136. 10.1016/j.coisb.2018.02.006

Wen, J. J., & Garg, N. J. (2018). Manganese Superoxide Dismutase Deficiency Exacerbates the Mitochondrial ROS Production and Oxidative Damage in Chagas Disease. PLOS Neglected Tropical Diseases, 12(7), e6687. 10.1371/journal.pntd.0006687

Wilson, D. F., & Matschinsky, F. M. (2021). Metabolic Homeostasis in Life as We Know It: Its Origin and Thermodynamic Basis. Frontiers in Physiology, 12, 658997. 10.3389/fphys.2021.658997

Winterbourn, C. C. (2013). The Biological Chemistry of Hydrogen Peroxide. In Hydrogen Peroxide and Cell Signaling, Part C: Vol. 528. Hydrogen Peroxide and Cell Signaling, Part C (pp. 3–25). Elsevier. 10.1016/B978-0-12-405881-1.00001-X

Yadav, D. K., Kumar, S., Choi, E.-H., Chaudhary, S., & Kim, M.-H. (2019). Molecular Dynamic Simulations of Oxidized Skin Lipid Bilayer and Permeability of Reactive Oxygen Species. Scientific Reports, 9(1), 4496. 10.1038/s41598-019-40913-y

Yan, L. L., & Zaher, H. S. (2019). How Do Cells Cope with RNA Damage and Its Consequences?. Journal of Biological Chemistry, 294(41), 15158–15171. 10.1074/jbc.REV119.006513

Yang, B. et al. (2019). Yin-Yang Mechanisms Regulating Lipid Peroxidation of Docosahexaenoic Acid and Arachidonic Acid in the Central Nervous System. Frontiers in Neurology, 10, 642. 10.3389/fneur.2019.00642

Yang, L. et al. (2019). Cellular Responses to Reactive Oxygen Species Are Predicted from Molecular Mechanisms. Proceedings of the National Academy of Sciences, 116(28), 14368–14373. 10.1073/pnas.1905039116

Yuan, Y., Ju, Y. S., Kim, Y., Li, J., Wang, Y., Yoon, C. J., Yang, Y., Martincorena, I., Creighton, C. J., Weinstein, J. N., Xu, Y., Han, L., Kim, H.-L., Nakagawa, H., Park, K., Campbell, P. J., Liang, H., PCAWG Consortium, Aaltonen, L. A.,…Von Mering, C. (2020). Comprehensive Molecular Characterization of Mitochondrial Genomes in Human Cancers. Nature Genetics, 52(3), 342–352. 10.1038/s41588-019-0557-x

Zhang, C.-Y., Parton, L. E., Ye, C. P., Krauss, S., Shen, R., Lin, C.-T., Porco, J. A., & Lowell, B. B. (2006). Genipin Inhibits UCP2-Mediated Proton Leak and Acutely Reverses Obesity- and High Glucose-Induced bn Cell Dysfunction in Isolated Pancreatic Islets. Cell Metabolism, 3(6), 417–427. 10.1016/j.cmet.2006.04.010

Zheng, M., Wang, X., Templeton, L. J., Smulski, D. R., LaRossa, R. A., & Storz, G. (2001). DNA Microarray-Mediated Transcriptional Profiling of the Escherichia Coli Response to Hydrogen Peroxide. Journal of Bacteriology, 183(15), 4562–4570. 10.1128/JB.183.15.4562-4570.2001

Zhong, J., Xiao, C., Gu, W., Du, G., Sun, X., He, Q.-Y., & Zhang, G. (2015). Transfer RNAs Mediate the Rapid Adaptation of Escherichia Coli to Oxidative Stress. PLOS Genetics, 11(6), e1005302. 10.1371/journal.pgen.1005302

Zhu, M., & Dai, X. (2019). Maintenance of Translational Elongation Rate Underlies the Survival of Escherichia Coli during Oxidative Stress. Nucleic Acids Research, 47(14), 7592–7604. 10.1093/nar/gkz467

